# Stochastic logistic models reproduce experimental time series of microbial communities

**DOI:** 10.1101/2020.01.31.928697

**Authors:** Lana Descheemaeker, Sophie de Buyl

## Abstract

We analyze properties of experimental microbial time series, from plankton and the human microbiome, and investigate whether stochastic generalized Lotka-Volterra models could reproduce those properties. We show that the noise term should be large and that it is a linear function of the species abundance, while the strength of the self-interactions should vary over multiple orders of magnitude. We stress the fact that all the observed stochastic properties can be obtained from a logistic model, i.e. without interactions, even the niche character of the experimental time series. Linear noise is associated with growth rate stochasticity, which is related to changes in the environment. This suggests that fluctuations in the sparsely sampled experimental time series are caused by extrinsic sources.

## Introduction

Microbial communities are found everywhere on earth, from oceans and soils to gastrointestinal tracts of animals, and play a key role in shaping ecological systems. Because of their importance for our health, human associated microbial communities have recently received a lot of attention. According to the latest estimates, for each human cell in our body we count one microbe (Sender et al., 2016). Dysbiosis in the gut microbiome is associated with many diseases from obesity, chronic inflammatory diseases, some types of cancer to autism spectrum disorder (Gilbert et al., 2016). It is therefore crucial to recognize what a healthy composition is, and if unbalanced, be able to shift the composition to a healthy state. This asks for an understanding of the ecological processes shaping the community and dynamical modeling.

Models of complex ecosystems typically focus on one specific aspect of the dynamics such as the stability of the community (May, 1972; Coyte et al., 2015; Levine et al., 2017; Grilli et al., 2017; Gavina et al., 2018; Gibbs et al.,2018),the neutrality (Fisher and Mehta, 2014; Washburne et al., 2016) or mechanisms leading to long-tailed rank abundance distributions (Solé et al., 2002; Brown et al., 2002; McGill et al., 2007; Matthews and Whittaker, 2015). Different types of dynamical models have been proposed. Generalized Lotka-Volterra (gLV) models describe the system at the population level and assume that the interactions between species dictate the community’s time evolution. Both deterministic and stochastic implementations exist for gLV models. Another approach consists in modeling the population at the individual level, which is referred to as agent-based models. Agent-based models include self-organized instability models (Solé et al., 2002) and the controversial neutral model of Hubbell (Hubbell, 2001; Rosindell et al., 2011) which assumes that all species are ecologically equivalent. A classification scheme that assesses the relative importance of different ecological processes from time series has been proposed in Faust et al. (2018). The scheme is based on testing for the temporal structure in the time series via an analysis of the noise color and neutrality. Applied to human stool microbiota time series, it tells us that stochastic gLV or self-organized instability models are more realistic. Here we will however only focus on stochastic gLV models. The reason for this is twofold. First, one can encompass the whole spectrum of ecosystems from neutral to niche with gLV models (Fisher and Mehta, 2014). Second, we aim at describing dense ecosystems and even though an individual based model might be more accurate, in the large number limit it will be captured by a Langevin approximation, *i.e*. by the stochastic gLV model.

Our goal is to compare time series generated by stochastic gLV models with experimental time series of microbial communities. We aim at capturing all observed properties mentioned above – the rank abundance profile, the noise color and the niche character - as well as the statistical properties of the differences between abundances at successive time points with one model. As is shown in Properties of experimental time series, the rank abundance profile is heavy-tailed, which means that few species are highly abundant and many species have low abundances. Despite the large differences in abundances, the ratios of abundances at successive time points and the noise color are independent of these abundances and although the fluctuations are large, the results of the neutrality tests indicate that the experimental time series are in the niche regime. To sum up, we seek for growth rates, interaction matrices, immigration rates and an implementation of the noise in stochastic gLV models to obtain the experimental characteristics.

We simulated time series using gLV equations. The interaction matrices are random as was introduced by May (1972). The growth rates are determined by the choice of the steady state, which is set to either equal abundances for all species or abundances according to the rank abundance profiles found for experimental data. For the noise, we consider different implementations corresponding to different sources of intrinsic and extrinsic noise.

Our analysis constraints the type of stochastic gLV models able to grasp the properties of experimental time series. First, we show that there is a correlation between the noise color and the product of the mean abundance and the self-interaction of a species. The noise color profile for such models will therefore depend on the steady state. This implies that imposing equal self-interaction strengths for all species which can be done to ensure stability (Fisher and Mehta, 2014; Gibson et al., 2016), is incompatible with properties of experimental time series. Second, from the differences between abundances at successive time points, we conclude that a model with mostly extrinsic (linear) noise agrees best with the experimental time series. Third, neutrality tests often result in the niche regime for time series generated by noninteracting species with noise. We therefore conclude that all stochastic properties of experimental time series are captured by a logistic model with large linear noise. However interactions are not incompatible with those properties. Our work raises caution for the interpretation of the character of the noise of time series, which goes along the lines of Carr et al. (2019) about the pitfalls of inferring interactions from covariance.

All codes are available online (see Additional files: Code).

## Properties of experimental time series

We study time series from different microbial systems: the human gut microbiome (David et al., 2014), marine plankton (Martin-Platero et al., 2018) and diverse body sites (handpalm, tongue, faecal) (Caporaso et al., 2011)(Figure 1A). A study of the different characteristics for a selection of this data is represented by Figure 1. The complete study of all time series can be found in the Supplemental information: Analysis of experimental data. We propose a detailed description of the properties of experimental time series. They fall essentially into two categories. The stability and rank abundance are tightly connected to the deterministic part of the equations while the differences between abundances at successive time points and noise color explain the stochastic behavior. The neutrality is more subtle and depends on the complete system.

**Figure 1.**
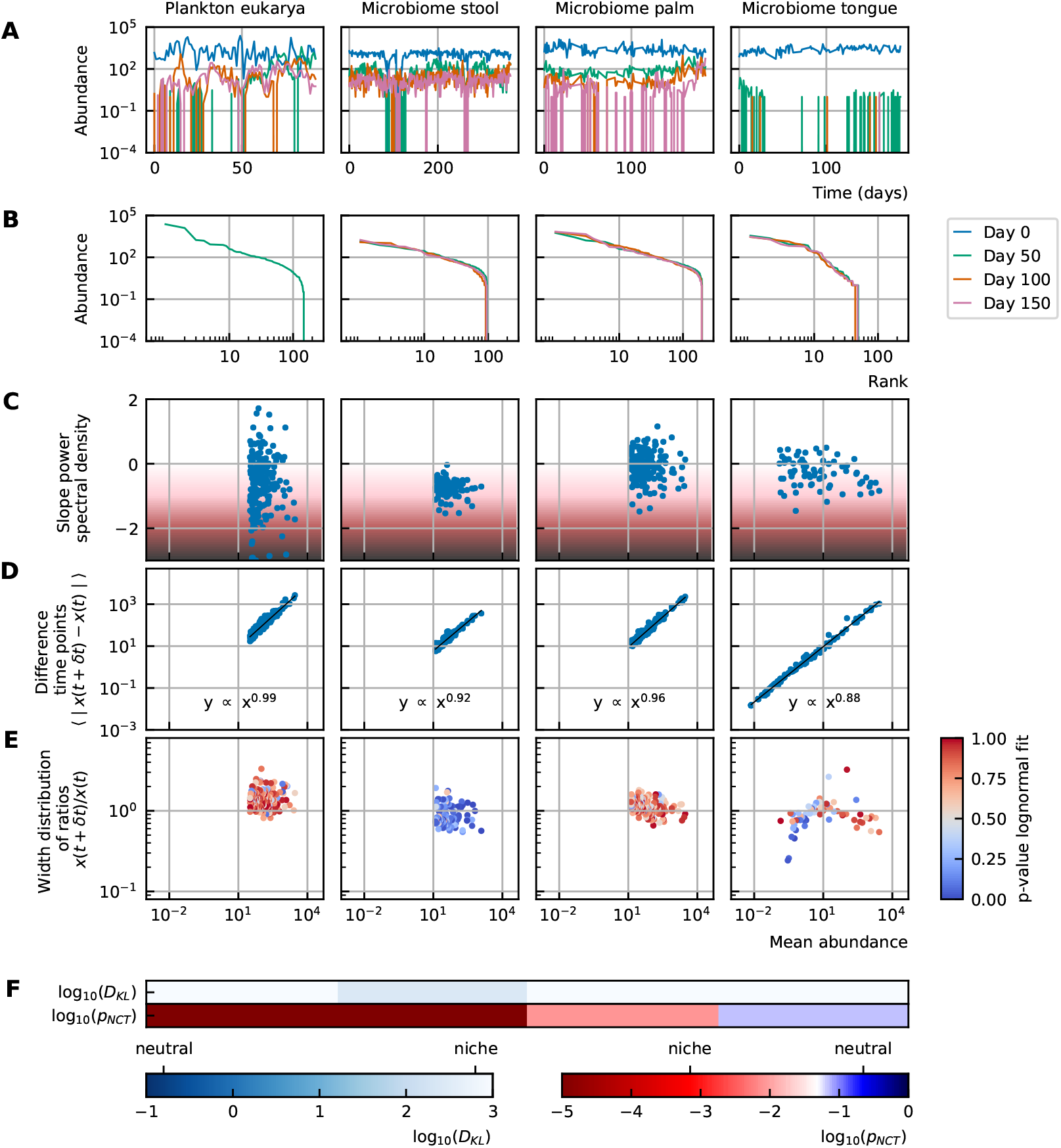
Characteristics of experimental data (A) Time series. (B) Rank abundance profile. The rank abundance is heavy-tailed and remains stable over time. (C) Noise color: No clear correlation between the slope of the power spectral density and the mean abundance of the species can be seen. The background colors represent the noise colors corresponding to the slope of the power spectral density (white, pink, brown, black). (D) Absolute difference between abundances at successive time points: There is a linear correspondence (in log-log scale) between the mean absolute difference between abundances at successive time points and the mean abundance of the species. Because the slope is almost one, this hints at the linear nature of the noise. (E) Width of the distribution of the ratios of abundances at successive time points: The width of the distribution of successive time points is large (order 1) and does not depend on the mean abundance of the species. Most of the species fit well a lognormal distribution: the p-values are high. (F) Neutrality: Both the Kullback-Leibler divergence (*D*_KL_) and the neutral covariance test (*p_NCT_*) indicate that most experimental time series are in the niche regime.

### The rank abundance can be described by a heavy-tailed distribution

The first aspect of community modeling that has been widely studied the last years is the stability of the steady states. Large random networks tend to be unstable (May, 1972). This problem is often solved by considering only weak interactions, sparse interaction matrices (May, 2001) or by introducing higher-order interactions (Grilli et al., 2017; Gavina et al., 2018; Sidhom and Galla, 2019). Although the stability of gLV models decreases with increasing number of participating species, the stability only depends on the interaction matrix and not on the abundances (Gibbs et al.,2018). The distribution of the steady state values is studied through the rank abundance.

The rank abundance distribution describes the commonness and rarity of all species. It can be represented by a rank abundance plot, in which the abundances of the species are given as a function of the rank of the species, where the rank is determined by sorting the species from high to low abundance. These curves can generally be described by power law, lognormal or logarithmic series functions (Limpert et al.,2001; McGill et al., 2007; Brown et al.,2002).

The rank abundance of the experimental time series can be described by a heavy-tailed distribution. This means that there are few common and many rare species. Rank abundances remain stable over time (Figure 1B).

### The differences between abundances at successive time points are large and linear with respect to the species abundance

Time series can be described by the differences between abundances at successive time points. We propose to focus on two specific representations of the information contained in those differences. First, we consider the mean absolute difference between abundances at successive time points 〈|*x*(*t* + *δt*) − *x*(*t*)|〉 as a function of the mean abundance 〈*x*(*t*)〉. For the experimental data, the relation between these variables is a monomial – this means that it is linear in a log-log representation (Figure 1D). The fact that the slope of this line is almost one hints at a linear nature of the noise.

Second, we examine the distribution of the ratios of the abundances at two successive time points *x*(*t* + *δt*)/*x*(*t*). The width of this distribution tells how large the fluctuations are. To measure this width we fit the distribution with a lognormal curve for which the mean is fixed to be one as the fluctuations occur around steady state. For most of the species of experimental data (except for the stool data), the fit of the distribution to a lognormal curve is good (high p-values, Figure 1E). Furthermore, we notice that the distribution is wide– in the order of1– and that the width does not depend on the mean abundance of the species (Figure 1 E). Examples of the fitted lognormal curve can be found in Supplemental information: Methods.

### The noise color is independent of the mean abundance of the species

The noise of a time series can be studied by considering the distribution of the frequencies of the fluctuations. The noise color is defined by the slope of a linear fit through the power spectral density. In this way, white, pink,brown and black noise correspond to slopes around 0, −1, −2 and −3 respectively. The more negative the slope is – this corresponds to darker noise – the more structure there is in the time series (Faust et al.,2018). More details about the estimation of the noise color can be found in the Supplemental information:Methods.

We notice that there is no correlation between the slope of the power spectral density (noise color) and the mean abundance of the species for experimental time series (Figure 1C).

### Experimental time series are in the niche regime

In neutral theory it is assumed that all species or individuals are functionally equivalent. It is challenging to test whether a given time series was generated by neutral or niche dynamics. We use two definitions of neutrality measures: the Kullback-Leibler divergence as used in Fisher and Mehta (2014) and the neutral covariance test as proposed by Washburne et al. (2016). The Kullback-Leibler divergence measures how different the multivariate distribution of species abundances is from a distribution constructed under the assumption of ecological neutrality. The idea of the neutral covariance test is to compare the time series with a Wright-Fisher process. A Wright-Fisher process is a continuous approximation of Hubbell’s neutral model for a large and finite community. In particular, it tests the invariance with respect to grouping. More about the implementation, interpretation and validity of these neutrality measures can be found in the Supplemental information: Methods.

Both neutrality measures, the Kullback-Leibler divergence and the neutral covariance test, indicate that most experimental time series are in the niche regime (Figure 1F).

## Modeling generalized Lotka-Volterra equations

In a microbial community different species interact because they compete for the same resources. Moreover, they produce byproducts that can affect the growth of other species. Depending on the nature of the byproducts, harmful, beneficial or even essential, the interaction strength will be either negative or positive. To describe the dynamics of interacting species, one can use the generalized Lotka-Volterra equations:

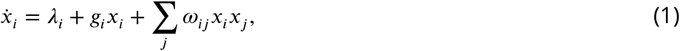

where *x_i_*, *λ_i_* and *g_i_* are the abundance, the immigration rate and the growth rate of species *i* respectively, and *ω_ij_* is the interaction coefficient that represents the effect of species *j* on species *i*. The diagonal elements of the interaction matrix *ω_ii_*, the so-called self-interactions, are negative to ensure stable steady states. The gLV equations only consider pairwise effects and no saturation terms or other higher-order terms. Due to this drawback, these models sometimes fail to predict microbial dynamics (Momeni et al., 2017; Levine et al., 2017). However, they are among the most simple models for interacting species and therefore widely studied and used. Noninteracting species can be described by the logistic model, which is a special case of the gLV model obtained by setting all off-diagonal elements of the interaction matrix to zero.

### Implementations of the noise

There exist two principle types of noise: intrinsic and extrinsic noise. *Extrinsic noise* arises due to external sources that can alter the values of the different variables: the immigration rate and growth rate can fluctuate in time through colonization of species or a changing flux of nutrients. These give rise to additive noise and linear multiplicative noise respectively. *Intrinsic noise* is due to the discrete nature of individual microbial cells. Thermal fluctuations at the molecular level determine the cell fitness of the individual cells. Therefore cell growth, cell division and cell death can be considered as stochastic Poisson processes. For large numbers of microbes these fluctuations will be averaged out.

We first consider the extrinsic noise. If the time series is calculated by *x_i_*(*t* + *dt*) = *x_i_*(*t*) + *dx_i_*(*t*), the implementation of the linear multiplicative noise is as follows,

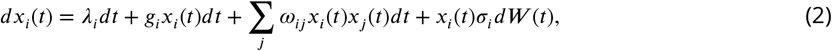

where *dW* is an infinitesimal element of a Brownian motion defined by a variance of 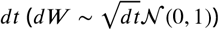. Changes in immigration rates of microbial species can be modeled with additive noise,

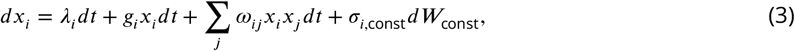

with 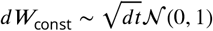. Our main motivation is to model the gut microbiome in the colon. Here, we ignore the immigration of species for two reasons. First, the number of microbes in the colon is orders of magnitude larger than the number of microbes in the other parts of the gut (Marteau et al., 2001; Gorbach et al., 1967), therefore the flux of incoming microbes in the colon is small. Second, we only consider systems around steady state for which we assume immigration does not play an important role. For perturbed systems, which are far from equilibrium, immigration rates can probably not be ignored. Ignoring immigration may be a too restrictive for other microbial systems such as the skin microbiome or plankton.

Intrinsic noise can be modeled by a noise term that scales with the square root of the species abundance (Walczak et al., 2012; Fisher and Mehta, 2014),

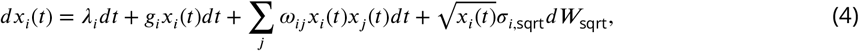

with *dW*_sqrt_ again an infinitesimal element of a Brownian motion defined by a variance of 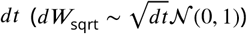. The size of this noise *σ*_*i*,sqrt_ is determined by the cell division (*g*^+^) and death rates (*g*^−^) separately which are in our model combined to one growth vector 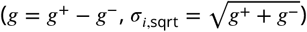, for large division and death rates the intrinsic noise will be larger.

To sum up, we focus on linear multiplicative noise because: (a) extrinsic noise is dominant as microbial communities contain a very large number of individuals and (b) we ignore immigration of individuals in our analysis.

We verified that our analysis is robust with respect to the multiple possibilities for the discretization of these models. We also compare our population level approach with individual based modeling approaches. Details can be found in the Supplemental information: Methods.

## Reproducing properties of experimental time series from stochastic generalized Lotka-Volterra models

### The rank abundance distribution can be imposed by fixing the growth rate

Random matrix models do typically not give rise to heavy-tailed abundance distributions. Neither is it known which properties of the interaction matrix and growth rates are required to obtain a realistic rank abundance distribution. We can however enforce a desired rank abundance artificially by solving the steady state of the gLV equations. For given steady state abundance vector 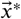 and interaction matrix *ω*, we impose the growth rate 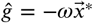. One model that results in heavy-tailed distributions is the self-organized instability model proposed by Solé et al. (2002).

### The noise color is determined by the mean abundance and the self-interaction of the species

To study the noise color, we first consider a model where the species are not interacting. The noise color is independent of the implementation of the noise, but depends on the product of the mean abundance and the self-interaction of the species (Figure 2A). For noninteracting species the growth rate equals the product of the self-interaction and the steady state abundance. Because we consider fluctuations around steady state, the mean and the steady state abundance are nearly equal and the *x*-axis of Figure 2A-C can be interpreted as the growth rate. Also, the strength of the noise does not change its color (Figure 2C). A parameter that is important for the noise color is the sampling rate: the higher the sampling frequency the more dark the noise becomes (Figure 2B). Darker noise corresponds to more structure in the time series. The more frequent the abundances are sampled the more details are visible and the underlying interactions become more visible. We conclude that the noise color is only dependent on the mean abundance, the self-interactions and the sampling rate. Figures of the dependence on the mean abundance and self-interaction separately can be found in the Supplemental information: Supporting results.

**Figure 2.**
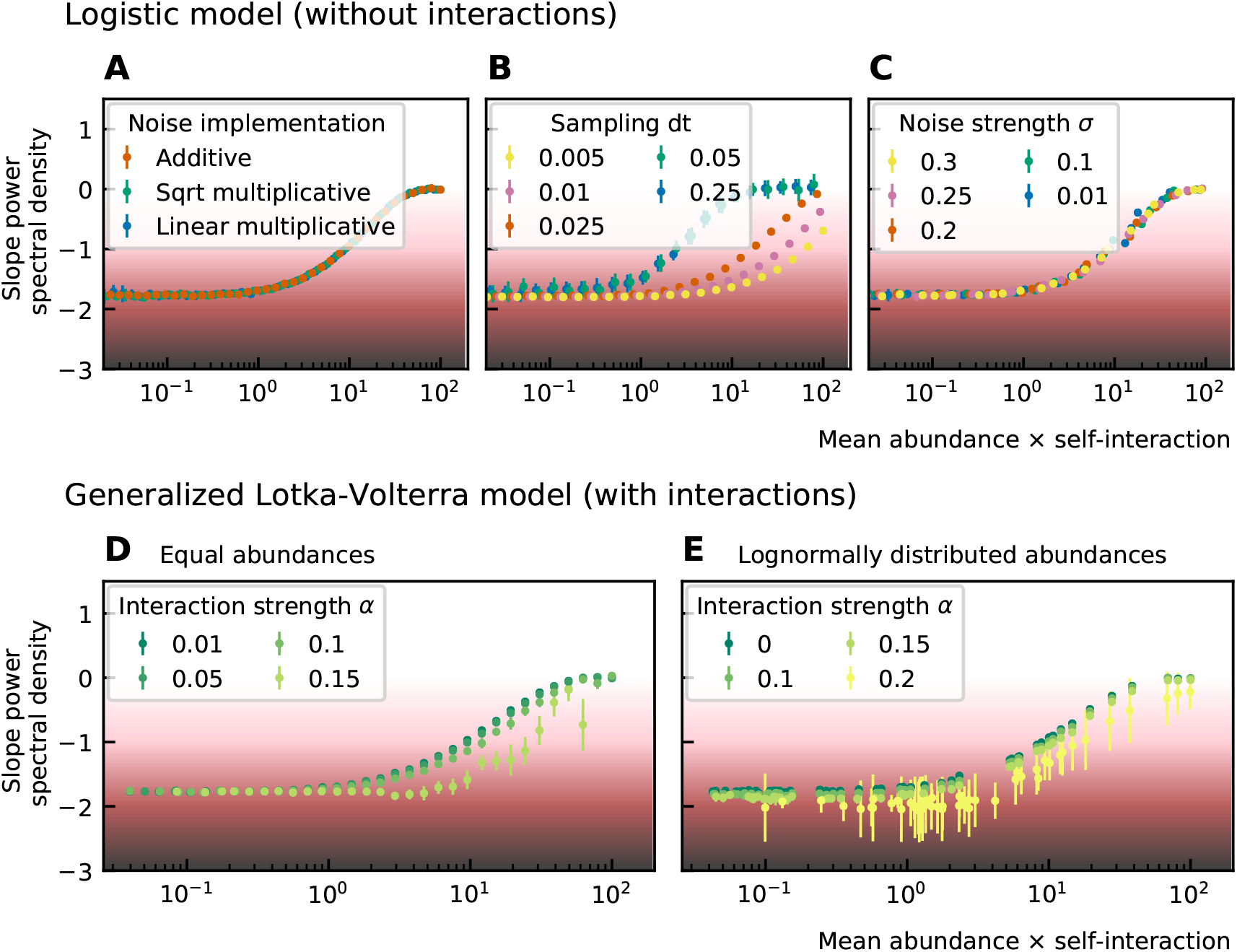
Noise color as a function of the mean abundance and self-interaction for stochastic logistic and gLV equations. The background colors represent the noise colors corresponding to the slope of the power spectral density (white, pink, brown, black). The mean abundance determines the noise color when there is no interaction, the implementation method (A) and the strength of the noise (C) have no influence. A smaller sampling time interval *δt*, which is equivalent to a higher sampling rate, makes the noise darker (B). For gLV models with interactions, larger interaction strengths make the noise colors darker for systems with equal abundances (D) as well as systems with heavy-tailed rank abundance distributions (E).

For interacting species increasing the strength of the interactions makes the color of the noise darker in the high mean abundance range (Figure 2D-E). Importantly, for interacting species with a lognormal rank abundance, the correlation between the noise color and mean abundance is preserved (Figure 2E). The data can be fit to obtain a bijective function between the product of the mean abundance and the self-interaction, and noise color. Assuming this model is correct, we can obtain an estimate for the self-interaction coefficients given the mean abundance and noise color by fixing the interaction strength. The uncertainty on the estimates is larger where the fitted curve is more flat (slopes of the power spectral density around −1.7 and 0), but many experimental values of the stool microbiome data lie in the pink region where the self-interaction can be estimated for this model.

### The implementation of the noise determines the correlation between the mean absolute increment 〈|*x*(*t* + *δt*) − *x*(*t*)|〉 and the mean abundance 〈*x*(*t*)〉

Next, we study the differences between abundances at successive time points. From the results of the noise color we can estimate the self-interaction for the dynamics of the experimental data. We use the rank abundance and the self-interaction inferred from noise color of the microbiome data of the human stool to perform simulations and calculate the characteristics of the distribution of differences between abundances at successive time points. We here assume that there are no interactions. More results for dynamics with interactions are in the Supplemental information: Supporting results. We first study the correlation between the mean absolute difference between abundances at successive time points 〈|*x*(*t* + *δt*) − *x*(*t*)|〉 and the mean abundance 〈*x*(*t*)〉. For linear multiplicative noise, the slope of the curve of the logarithm of the mean absolute difference between abundances at successive time points log_10_(〈|*x*(*t* + *δt*) − *x*(*t*)|〉) as a function of the logarithm of the mean abundance log_10_(〈*x*(*t*)〉) is one. For multiplicative noise that scales with the square root of the abundance, the slope is around 0.66 and for additive noise, the slope is zero. By combining both linear noise and noise that scales with the square root of the abundance slopes with values between 0.6 and 1 can be obtained (Figure 3A). The slopes of experimental data range between 0.84 and 0.99, we therefore conclude that linear noise is a relatively good approximation to perform stochastic modeling of microbial communities.

**Figure 3.**
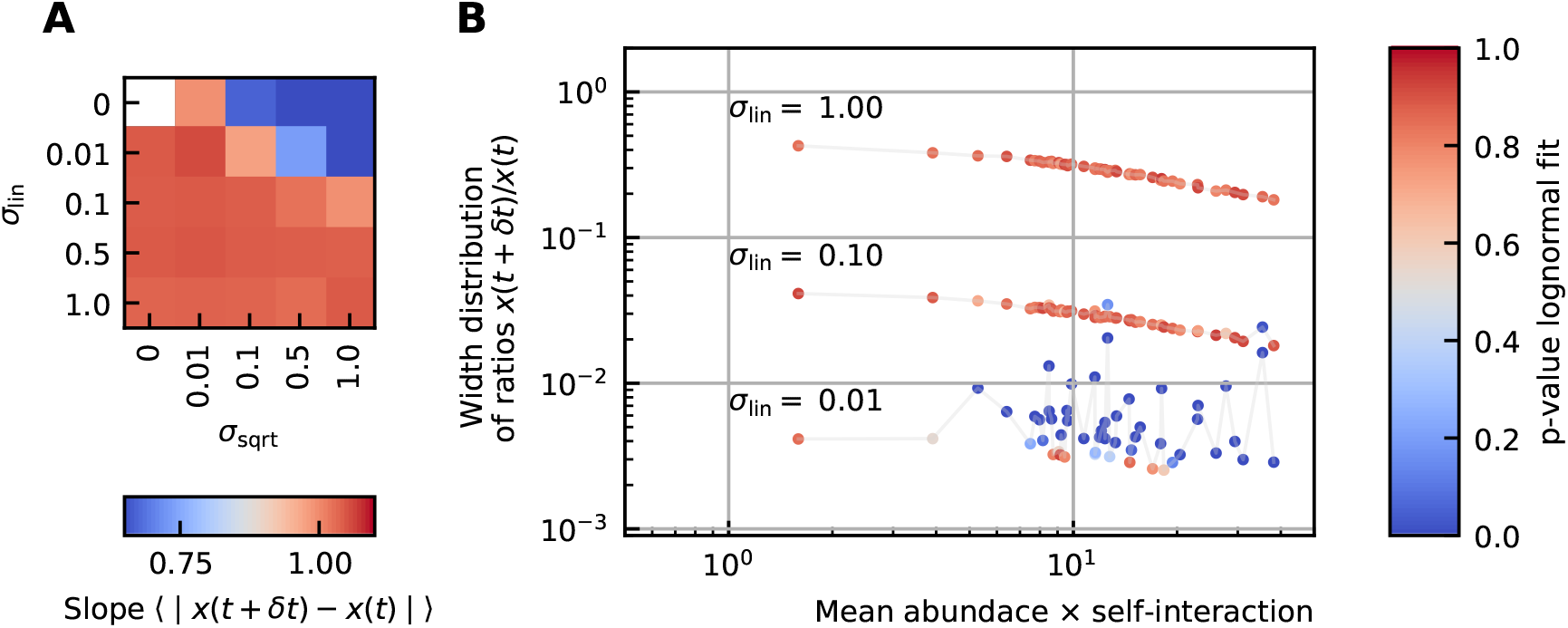
(A) The slope of the logarithm of the mean absolute difference between abundances at successive time points as a function of the logarithm of the mean abundance for different strengths of the linear noise (*σ*_lin_) and multiplicative noise that scales with the square root of the abundances (*σ*_sqrt_). The slope ranges from 0.66 for noise that scales with the square root to 1 for linear noise. (B) The width of the distribution of the ratios of abundances at successive time points increases for increasing strength of the noise. For sufficiently strong noise the distribution is well fitted by a lognormal function (high p-values).

### The strength of the noise determines the width of the distribution of ratios *x*(*t* + *δt*)/*x*(*t*)

Next we examine the distribution of the ratios of abundances at successive time points. As expected, for significant noise this distribution can be approximated by a lognormal curve and the width of the distribution becomes larger for increasing noise strength (Figure 3B). In order to have widths that are in the same order of the ones of the experimental data, the noise must be sufficiently strong. Another way of increasing the width is through interactions, this effect is only moderate. These results are presented in the Supplemental information: Supporting results.

### Stochastic logistic models capture the properties of experimental time series

By using all previous results and imposing the steady state of experimental data, we find that it is possible to generate time series with identical characteristics as seen in experimental time series (Figure 4). Furthermore these time series can be generated without introducing any interaction between the different species, but their neutrality measures can still be in the niche regime (Figure 4C). Out of 100 simulations 62 had a p-value smaller than 0.05 for the neutral covariance test which means they are in the niche regime. The colors of the noise fix the self-interaction values (Figure 4D), next the rank abundance distribution is imposed by calculating the growth vector *ĝ* (Figure 4B). The slope of the curve of the mean absolute difference between abundances at successive time points as a function of the mean abundance is one by using linear multiplicative noise (Figure 4E) and the width of the fluctuations is tuned by choosing a large noise size *σ* (Figure 4F). Similar results can be obtained for models with interactions (see Supplemental information: Supporting results), but we want to stress that interactions are not needed to reproduce the properties of experimental time series.

**Figure 4.**
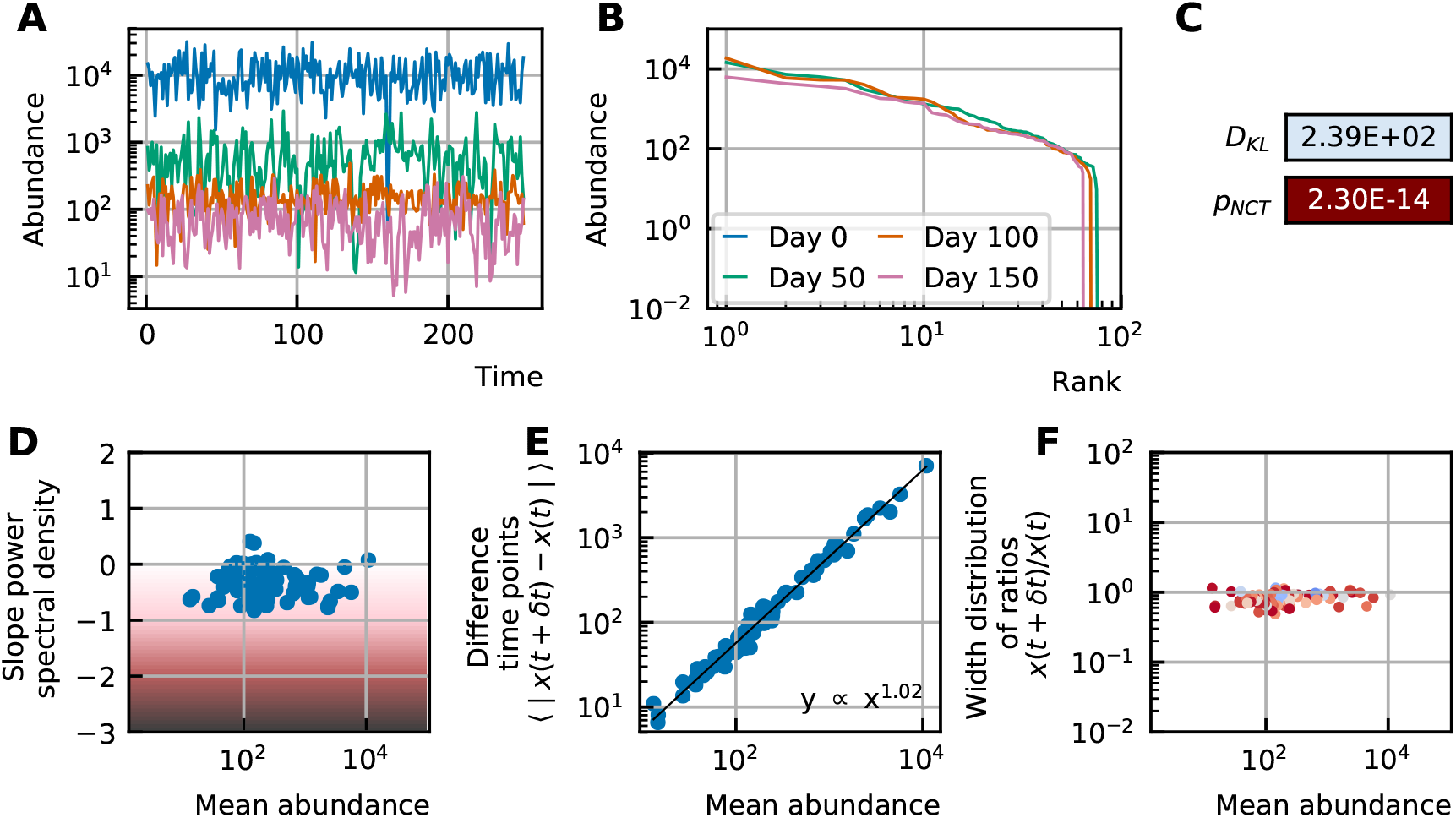
A stochastic logistic model is able to reproduce the different characteristics of the noise: (A) Time series. (B) A heavy-tailed rank abundance that remains stable overtime. (C) Results of the neutrality test in the niche regime. (D) Noise color in the white-pink region with no dependence on the mean abundance. (E)The slope of the mean absolute difference between abundances at successive time points is around 1. (F) The width of the distribution of the ratios of abundances at successive time points is in the order of 1 and independent of the mean abundance.

## Discussion

Recent research has focused on different aspects of experimental time series of microbial dynamics, in particular the rank abundance distribution, the noise color, the stability and neutrality. Within the framework of stochastic generalized Lotka-Volterra models, we studied the influence of growth rates, interactions between species and the different sources of stochasticity on the observed characteristics of the noise and on neutrality. Our observations are:

- Even when we consider the case of no interactions between species, the result of the neutrality test on the time series is often niche. We should therefore be careful in the interpretation of the results of neutrality tests.
- The noise color depends on the product of the self-interaction and the mean abundance, which for noninteracting species reduces to a dependence on the growth rate. Assuming the model can be used for microbial communities, the self-interaction coefficients can be estimated given the mean abundance and noise color. For the experimental time series (plankton, gut and human microbiome) the self-interaction strengths range over several orders of magnitude. The convention of equalling all self-interactions to −1 used in several studies (Fisher and Mehta, 2014; Gibson et al., 2016), cannot be adopted for stochastic models of communities with a heavy-tailed rank abundance distribution.
- The exponent of the mean absolute differences between abundances at successive time points with respect to the mean abundances is slightly smaller than one for experimental time series. Linear multiplicative noise results in a value of one, square root noise results in lower values (0.6). A mix of linear and square root noise can result in slopes with intermediate values.
- A large multiplicative linear noise is in agreement with both the distribution of the ratios of abundances at successive time points and the relation between the differences between abundances at successive time points and mean values.

To conclude, characteristics of experimental time series, from plankton to gut microbiota, can be reproduced by stochastic logistic models with a dominant linear noise. We expect however that for higher sampling rates, modeling the interactions between microbes would be necessary to explain the properties of the time series. For gut microbial time series, the system is sampled only once a day and therefore dominated by the noise in the growth terms corresponding to a linear noise.

Predictive models for microbial communities dynamics will certainly require a more in-depth description of the system. Nutrients and spatial distribution of microbes should play a role to dictate the evolution of the community, as well as the interaction with the environment. Synthetic microbial communities are currently being developed and will hopefully provide a more comprehensive view on the complexity of microbial communities (Vrancken et al., 2019).

## Acknowledgments

We thank Karoline Faust for interesting discussions, Pankaj Mehta for clarifications about his work on the niche-neutral transition and Stefan Vet and Lendert Gelens for careful reading of the manuscript.

## Additional files

### Supplemental information

#### Methods

Detailed description of the methods: neutrality measures, distribution of the ratios of abundances at successive time points, implementation and interpretation of the noise, noise color measurement, discretizations of stochastic models.

#### Analysis of experimental data

Analysis of all experimental data. Rank abundance, distribution of the differences between abundance at successive time points, neutrality tests and noise color.

#### Supporting results

Supporting results.

#### Code

All python codes to perform time series simulations, analysis and make all different figures of the main paper and supplement are available at https://github.com/lanadescheemaeker/logistic_models.

