## Supplemental material for "Stochastic logistic models reproduce experimental time series of microbial communities"

### Methods

#### Neutrality measures

There is no consensus on the definition of neutrality. In general, ecosystems are considered neutral if the dominating cause of fluctuations are random birth and death processes and not fitness advantages of species.

Different neutrality measures focus on different aspects of neutrality. The Kullback-Leibler divergence verifies whether all species are equal (equal abundances and equal covariances). The neutrality covariance test studies the grouping invariance of species in time series.

Given two distributions $P$ and $Q$ the *Kullback-Leibler divergence* is defined as

$$D_{KL}(P|Q) = E_P \left[ \ln \frac{P}{Q} \right] \tag{1}$$

where $E_P$ is the expectation value using the probabilities of distribution $P$. The density function of a multivariate Gaussian distribution is

$$P(x) = \frac{1}{(2\pi)^{N/2}\sqrt{\det K}} \exp\left(-\frac{1}{2}(x-\mu)^T K^{-1}(x-\mu)\right) \tag{2}$$

where $\mu$ and $K$ are the mean and covariance matrix of the distribution respectively. The Kullback-Leibler divergence for two multivariate Gaussian distributions in $\mathbb{R}^n$ is (Duchi, 2007)

$$D_{KL}(P|Q) = \frac{1}{2}\left(\ln\frac{\det K_Q}{\det K_P} - n + \mathrm{Tr}\left(K_Q^{-1}K_P\right) + (\mu_Q - \mu_P)^T K_Q^{-1}(\mu_Q - \mu_P)\right). \tag{3}$$

For every time series, we can calculate the $\mu$ and $K$ and we define values $\mu_N$ and $K_N$ for a corresponding neutral time series in which all species are equal (Fisher and Mehta, 2014). The distance to neutrality $D_{KL}(P|P_N)$ can thus be calculated by computing the probability distribution of the original time series $P$ and the associated neutral distribution $P_N$ with mean values $\mu_N = S^{-1}\sum_{i=1}^{S}\mu_i$ and $K_{P,ii} = S^{-1}\sum_{i=1}^{S}K_{ii}$ and $K_{P,ij} = S^{-1}(S-1)^{-1}\sum_{i=1}^{S}\sum_{j=1,i\neq j}^{S}K_{ij}$ with $S$ the number of species.

The *neutral covariance test* was designed by Washburne et al. (2016). We used a python translation of the code developed by this author.

#### Validity of neutrality measures

One way of generating a neutral time series is considering a lattice of $N$ individuals on which every time step one individual is replaced by another. Each individual of the lattice has an equal probability of being replaced (probability is $N^{-1}$). The disappearance of the first species can be interpreted as the result of either death or emigration. The replacing individual is either the result of immigration or growth. The probability of immigration depends on the immigration rate $\lambda$ ($0 \leq \lambda \leq 1$). In case of an immigration event, all species of the external species pool $S$ have an equal probability of immigrating. The probability of a growth event is thus given by the remaining $1-\lambda$. In case of growth, every individual has an equal probability of growing. Time series generated in this way depend on three variables: the length of the simulation time $T$, the immigration probability $\lambda$ and the number of individuals $N$. We study the effect of these three variables on both neutrality measures (Figure 1).

As discussed in Wennekes et al. (2012), the neutral-niche debate is a scale problem. Looking at a process from close-by and describing all the details, results in the niche regime. Dynamics of large systems where the details are not modeled appear to be dominated by random processes. The questions are: at which scale is one studying the process and which of the deterministic or stochastic processes is dominating? We can see this in the upper row of Figure 1A. Larger immigration gives rise to a neutral regime, however for short time series and large communities one will only see the "details" because the time series is not yet in steady state. Consequently this results in niche values. In the absence of immigration, the "winner-take-all" principle holds. During a transient period all species but one go extinct in a stochastic manner. In the end, the steady state consists of only one species surviving. The duration of the transient depends on the size of the community. This is why the time series are in the niche regime with respect to grouping in the absence of immigration when the time series is long enough (Figure 1B). For small immigration and small communities,

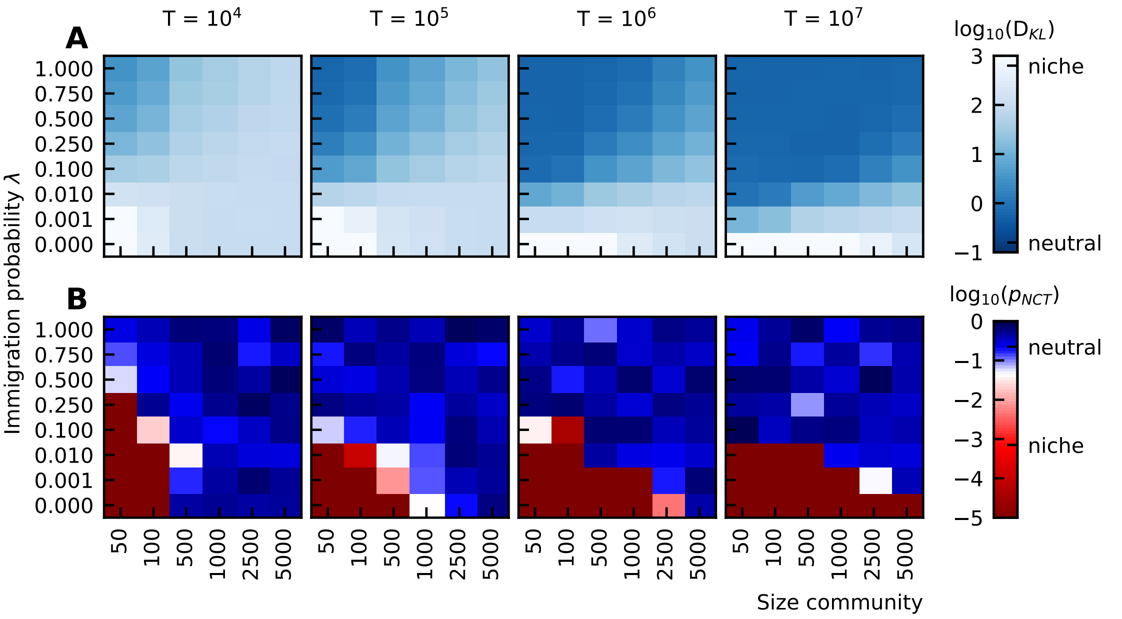

**Figure 1.** Neutrality tests, (A) Kullback-Leibler divergence and (B) the p-value of the neutral covariance test. We expect to see the neutral regime for all time series because they are generated by random birth, death and immigration events. For short time series and large size of the community, only the details are seen and the result is niche. For small immigration, there is a "winner-take-all" effect, for which the neutrality covariance test, which tests for invariance with respect to grouping gives a niche result.

the "winner" cannot take over due to the immigration of other species, but the most fit species dominate the community one by one in a random order. For short time series, there are only a few changes of dominating species and this apparent structure in the time series gives rise to a niche regime (Figure 1B). For longer time series however, there are many random changes of the dominating species, which results in a neutral character (Figure 1B).

#### Width distribution of ratios of abundances at successive time points

To study the jumps that the species abundances make over time, we consider the distribution of the ratios of abundances at successive time points $x(t + \delta t)/x(t)$. This distribution can be fitted to a lognormal function:

$$f(x) = \frac{1}{\sqrt{2\pi} s x} \exp\left(-\frac{\ln^2(x)}{2s^2}\right). \tag{4}$$

We impose the constraint that the median of this distribution is one because we assume that the time series fluctuates around steady state and that there will be as many steps for which the abundance is increasing as there are steps for which the abundance is decreasing. We define the shape parameter $s$ as the width of the step distribution. Some examples of fits are shown in Figure 2. The goodness of the fit is characterized by the p-value, a higher p-value corresponds to a better fit of the distribution to a lognormal function. The p-value is represented by the color of the fitted line.

#### Implementation and interpretation of the noise

Noise is introduced in dynamical systems by different processes. In simulations of microbial communities at the species level we distinguish intrinsic and extrinsic noise. Intrinsic noise is a result of the stochastic nature of cell division and cell death together with the discreteness of individuals. Extrinsic noise is caused by environmental fluctuations, e.g. changing nutrients, pH, temperature. To implement this extrinsic noise in a system of generalized Lotka-Volterra equations, the changing environments are often interpreted in changing parameters such as the growth rate. Because this parameter is multiplied by the species abundance, the

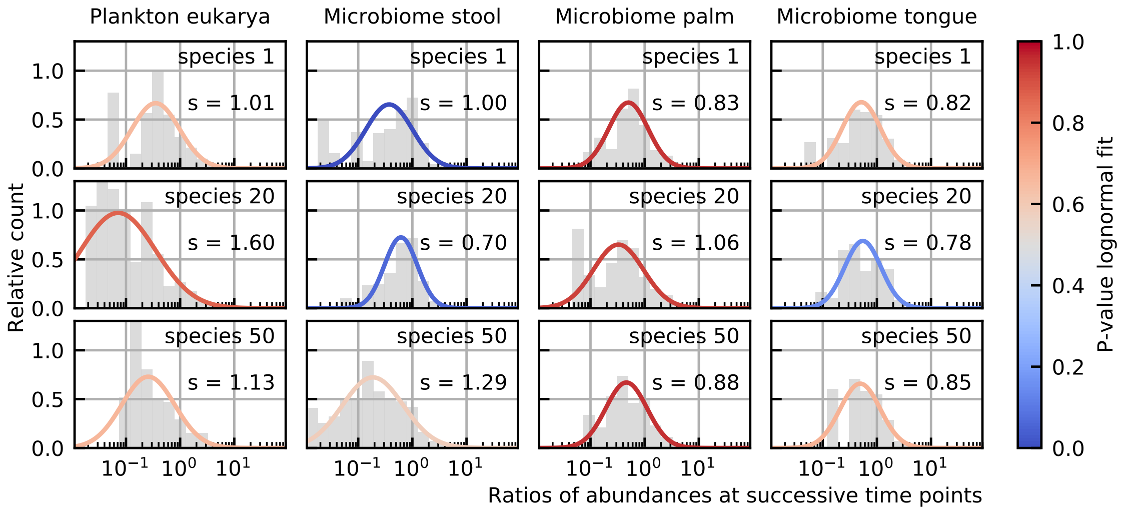

**Figure 2.** A lognormal function is fitted to the ratios of abundances at successive time points $x(t+\delta t)/x(t)$ for different species. The species number denotes its rank. We interpret the shape parameter $s$ as a measure for the width of the steps. Here we see that this width $s$ is of the order of 1 for all species. The goodness of the fit (p-value) is represented by the color of the fitted line. For most species the fit is good (high p-values).

resulting noise is linear. The remaining parameters, inter- and intra-specific interactions can also change depending on the environment. The formulation of this noise is more subtle (used in Zhu and Yin (2009)).

To derive the form of intrinsic noise in generalized Lotka-Volterra equations, we can consider every species abundance making a random walk in one dimension. The average displacement is zero and the variance of displacement is the sum of the rate of growth (jumping to the right) and the rate of death (jumping to the left). For the generalized Lotka-Volterra equations this results in a noise term

$$\langle n_i(t)n_i(t')\rangle = (\text{growth rate}_i + \text{death rate}_i)x_i = (f(g_i) + h(\omega, \vec{x}))x_i\delta(t-t'), \quad (5)$$

with $\omega$ the interaction matrix and where functions $f$ and $h$ decouple the growth and death terms. In the generalized Lotka-Volterra model no difference is made between negative interactions as a result from slowing down the growth rate or increasing the death rate, only the resulting net rates are used. This distinction must however be made to implement the intrinsic noise for gLV. In our analysis, we use the simpler logistic models where the resulting variance of the noise is proportional to the square root of the abundance $\sqrt{x}$. One must be careful not to use this noise for values that are smaller than 1, because this derivation relies on Poisson statistics which is defined for integer numbers.

### Noise color

The color of the noise in a time series is determined by the slope of the power spectral density in log-log scale. This slope can be determined by a linear fit through the spectrum. A different technique has been put forward to estimate the slope of the spectral density by Faust et al. (2018). There it is argued that the power spectral density does not have a constant slope and that therefore a nonlinear curve must be fitted. They choose for a spline fit and consider the minimal value of its derivative as the value for the noise color. Because the minimal value of the slope of the fit is taken, the noise color tends to be darker when using this technique. For our time series however we see that the spline fit only deviates from the linear fit for low frequencies (Figure 3). We ignore the low frequencies for fitting because of the windowing effect. Therefore, we opt for a linear fit after omitting the values for low frequencies (one order of magnitude of the lowest frequencies).

The correspondence between the colors and slopes is here:

As explained in the supplemental material of Faust et al. (2018), noise color is related to the autocorrelation of the time series. The lighter the noise color is, the steeper the autocorrelation function is at time delay

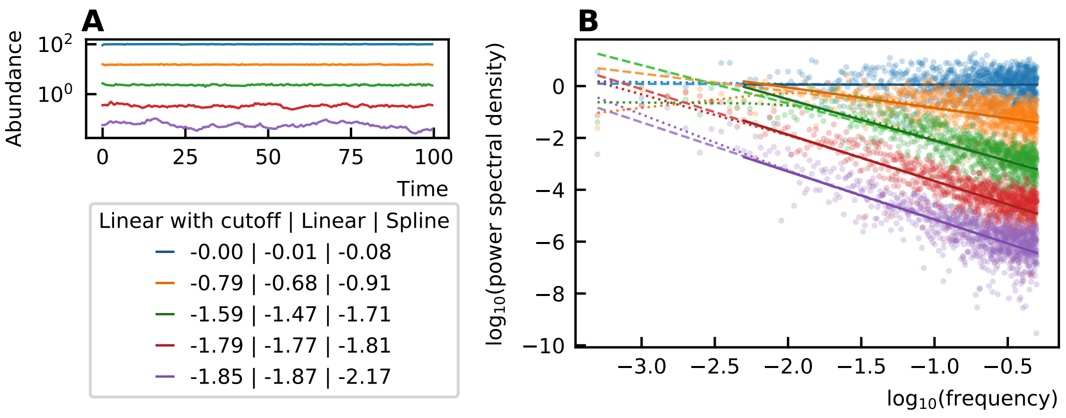

**Figure 3.** The noise color of time series (A) is determined by the slope of the power spectral density (B). This slope can be measured through a linear fit of all values (dashed), a linear fit through the higher frequency range (solid line) or by performing a spline fit (dotted). A linear fit through all frequencies can be influenced by the windowing effect for low frequencies and the spline fit can make the slope steeper at the low frequencies and result in a darker noise as can be seen for the purple curves. The values of the noise color determined by the different techniques are given in the legend. Therefore, in our work, we opt for the linear fit with a cutoff for low frequencies.

| Slope | Color |
|---|---|
| 0 | white |
| -1 | pink |
| -2 | brown |
| -3 | black |

$\delta = 0$. For dark noise, the autocorrelation decreases more slowly and the time series is said to contain more structure. An illustration of this is given by Figure 4.

#### Discretizations of stochastic models with linear noise

The implementation of the linear multiplicative noise is as follows,

$$dx_i = \lambda_i dt + g_i x_i dt + \sum_j \omega_{ij} x_i x_j dt + x_i \sigma_i dW, \tag{6}$$

with $dW$ an infinitesimal element of a Brownian motion which is defined by a variance of $dt$ ($dW \sim \sqrt{dt}\mathcal{N}(0,1)$).

Because of its nonlinear nature, generalized Lotka-Volterra equations cannot be solved analytically. It is therefore necessary to perform simulations of the stochastic differential equations.

When we consider a linear multiplicative noise, we must first choose a discretization of the deterministic differential equations. Two possibilities can be found in literature. The first one is the Ricker implementation and the second one which we are here calling the Langevin implementation.

#### Langevin discretization

To solve the stochastic equations numerically, we can use the Euler-Maruyama method. Here the discretization becomes

$$x_i(t + \delta t) = x_i(t) + g_i x_i(t)\delta t + \omega_{ij} x_i(t) x_j(t)\delta t + \sigma \delta W x_i(t). \tag{7}$$

Due to the choice of discretization $x$ can become zero or negative. Because this is the most straightforward discretization of the Langevin equation, we call it the Langevin discretization.

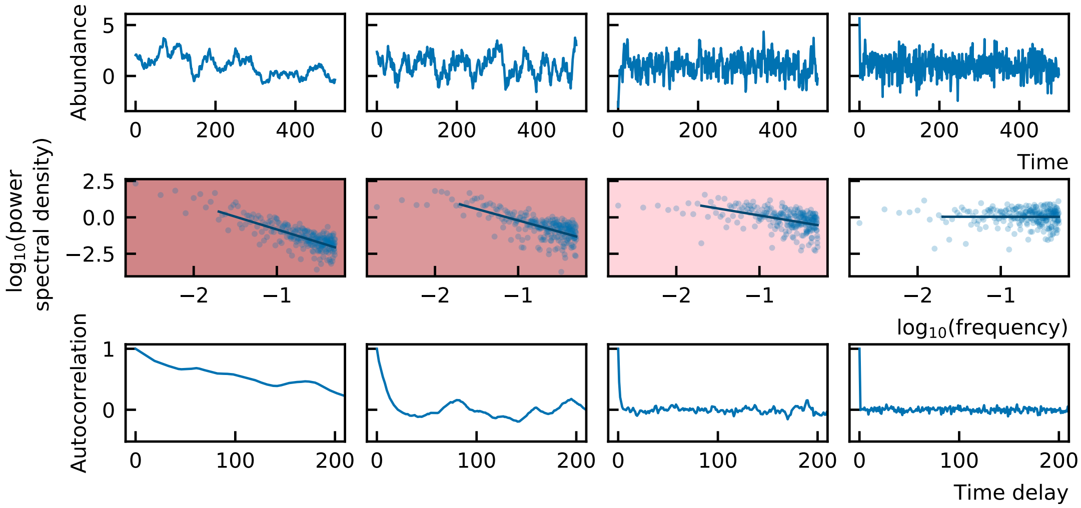

**Figure 4.** Different stochastic time series (first row) with the corresponding power spectral density (second row) and autocorrelation functions (last row). The background color of the power spectral density corresponds to the color of the noise. When there is more structure present in the time series, the autocorrelation function is less steep at time delay $\delta = 0$ and the noise is darker.

### Ricker discretization

For this implementation, we assume there is no migration ($\lambda_i = 0$). First, we rewrite Equation 6 as the derivative of the species abundance divided by the species abundance itself, such that we can replace the latter by a derivative of the logarithm of the species abundance:

$$\frac{\dot{x}_i}{x_i} = g_i + \sum_j \omega_{ij} x_j + \sigma \frac{dW}{dt}, \tag{8}$$

$$\dot{\ln x_i} = g_i + \sum_j \omega_{ij} x_j + \sigma \frac{dW}{dt}. \tag{9}$$

When we divided by the species abundance, we made the assumption that $x_i$ can never become zero (or negative for the solution is continuous). This would be true in the theoretically continuous framework and in the absence of noise, but this is not what is observed in experimental time series where the abundances are discrete numbers and can become zero in the event of extinction.

Using the assumption $\delta t \ll 1$, we obtain

$$\ln x_{t+\delta t} - \ln x_t = \delta t(\omega x + g) + \sigma \delta W, \tag{10}$$

$$x_{t+\delta t} = x_t \exp \delta t(\omega x + g) \exp \sigma \delta W \tag{11}$$

The $\delta W$ are normally distributed random elements ($\delta W \sim \sqrt{\delta t}\mathcal{N}(0,1)$).

Due to the choice of discretization, $x_i$ can never become zero or negative, which is fundamentally different from the previous implementation.

### Arato discretization

Using Ito calculus (see Box 1 for a short introduction) a third discretization can be obtained:

$$x_i(t) = x_i(0) \exp \left( g_i t - \frac{\sigma_i^2 t}{2} - \sum_{j=1}^{n} \omega_{ij} \int_0^t x_j(s) ds + \sigma W_i(t) \right) \tag{12}$$

as calculated in Arató (2003).

In the limit of small noise or small timesteps, the Langevin, Ricker and Arato discretizations are equal (Figure 5A). For large noise a clear difference can be seen for both implementations (Figure 5B).

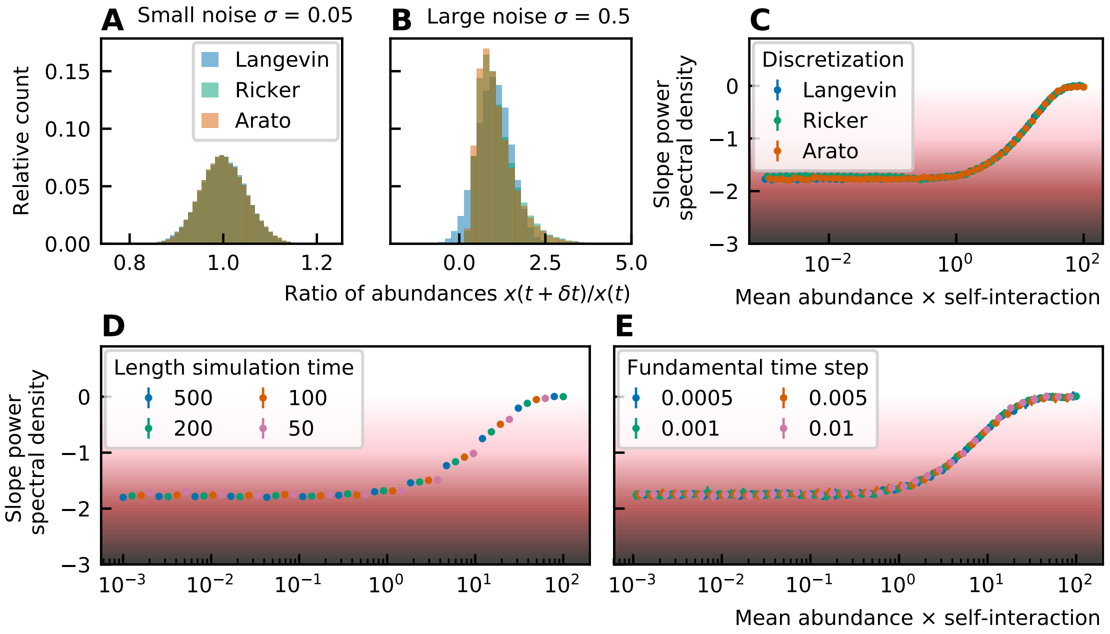

**Figure 5.** Distributions of $x_{t+\delta t}$ for $x_t = 1$ for three different discretizations: Langevin $x_{t+\delta t} = x_t + x_t \sigma \delta W$, Ricker $x_{t+\delta t} = x_t \exp(\sigma \delta W)$ and Arato $x_{t+\delta t} = x_t \exp\left(\frac{-\sigma^2}{2}\delta t + \sigma \delta W\right)$ with $\delta W \sim \mathcal{N}(0,1)$ and $\delta t = \sigma^2$ for small noise $\sigma = 0.05$ (A) and large noise $\sigma = 0.5$ (B). For small noise all implementations result in the same abundance distribution, for large noise the distributions are different. The Langevin implementation allows abundances to become zero and negative (which will be equalled to zero). The abundances of the Ricker and Arato implementation never become zero. The noise color does not depend on the particular implementation: it is independent of the discretization (C), length of the time series (D) and fundamental time step (E).

---

The parameters of the implementation, such as the discretization, length of the simulation and fundamental time step are not influencing the results of the noise color (Figure 5C-E).

#### Discretizations of stochastic models with non-linear noise

In the main manuscript, we discussed that we chose for a noise term that is linear in the species abundance, because this respresents best the external noise on growth and death of the species. Intrinsic noise (due to discreteness) could be represented by a term that is proportional to the square root of the species abundance (sqrt multiplicative noise) (Walczak et al., 2012). In this case the discretization is:

$$x_i(t+\delta t) = x_i(t) + g_i x_i(t)\delta t + \omega_{ij} x_i(t) x_j(t) \delta t + \sigma \delta W \sqrt{x_i(t)}. \quad (16)$$

We also considered a constant noise term (additive noise), which represents noise due to stochastic immigration and emigration of species. The corresponding discretization is:

$$x_i(t+\delta t) = x_i(t) + g_i x_i(t)\delta t + \omega_{ij} x_i(t) x_j(t) \delta t + \sigma \delta W. \quad (17)$$

In the main paper it is shown that the source of the noise, linear multiplicative, square root multiplicative or additive, has no influence on the relation between the mean abundance, self-interaction and noise color.

### Analysis of experimental data

We studied time series of different microbial communities. References for all of the time series can be found in Table 1.

#### Rank abundance

Rank abundances of all studied time series can be found in Figure 6. In nature, rank abundances are often power law, lognormal or logarithmic series distributions. The rank abundance does not fit one of these distributions exactly, but it is likewise heavy-tailed (Figure 7).

### Box 1. Basics of Ito calculus

A Brownian motion or Wiener process is described by a probability distribution over the set of continuous functions $B : \mathbb{R}_{\geq 0} \to \mathbb{R}$ which is defined by three characteristics:

1. $P(B(0) = 0) = 1$, the motion starts at the origin,
2. the motion is stationary: $\forall 0 \leq s \leq t : B(t) - B(s) \sim \mathcal{N}(0, t - s)$,
3. the increments are independent: if intervals $[s_i, t_i]$ are not overlapping than $B(t_i) - B(s_i)$ are independent.

An important feature of such a motion is the quadratic variation, which states that the expectation value of $dW^2$ is $dt$. In a regular derivative, the square and other higher order terms of the infinitesimal $dt$ of the Taylor expansion are ignored. Due to the quadratic variation, the square term of $dW$ cannot be ignored and Ito's lemma states that for a stochastic process $X_t$:

$$dX_t = \mu dt + \sigma dW_t, \tag{13}$$

and for $f$ a smooth function, we have that

$$df(t, X_t) = \left( \frac{\partial f}{\partial t} + \mu \frac{\partial f}{\partial x} + \frac{1}{2}\sigma^2 \frac{\partial^2 f}{\partial x^2} \right) dt + \frac{\partial f}{\partial x} dW_t, \tag{14}$$

and as a consequence,

$$d(\ln x_i) = \frac{dx_i}{x_i} - \frac{(dx_i)^2}{2x_i^2}. \tag{15}$$

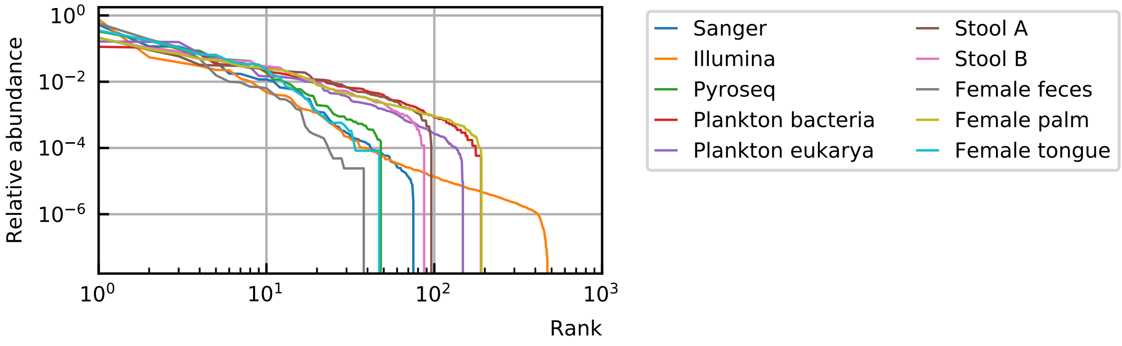

**Figure 6.** The rank abundance curve of experimental measurements of microbial communities, from human microbiomes to plankton, is heavy-tailed.

**Table 1.** References for all time series and compositions of microbial communities.

| Label | Label in Figure 1 of the main article | Source |
|---|---|---|
| Stool A | Microbiome stool | Subject A gut of David et al. (2014) |
| Stool B | | Subject B gut of David et al. (2014) |
| Plankton bacteria | | Bacteria relative abundance of Martin-Platero et al. (2018) |
| Plankton eukarya | Plankton eukarya | Eukaryota relative abundance of Martin-Platero et al. (2018) |
| Female feces | | Feces microbiome of subject F4 at the genus level (L6) Caporaso et al. (2011) |
| Male feces | | Feces microbiome of subject M3 at the genus level (L6) Caporaso et al. (2011) |
| Female left palm | Microbiome palm | Left palm microbiome of subject F4 at the genus level (L6) Caporaso et al. (2011) |
| Male left palm | | Left palm microbiome of subject M3 at the genus level (L6) Caporaso et al. (2011) |
| Female right palm | | Right palm microbiome of subject F4 at the genus level (L6) Caporaso et al. (2011) |
| Male right palm | | Right palm microbiome of subject M3 at the genus level (L6) Caporaso et al. (2011) |
| Female tongue | Microbiome tongue | Tongue microbiome of subject F4 at the genus level ((L6) Caporaso et al. (2011) |
| Male tongue | | Tongue microbiome of subject M3 at the genus level (L6) Caporaso et al. (2011) |
| Sanger | | Feces composition of DA-AD-1 of MetaHIT Consortium (additional members) et al. (2011) |
| Illumina | | Feces composition of MH0001 of MetaHIT Consortium (additional members) et al. (2011) (original data MetaHIT Consortium et al. (2010)) |
| Pyroseq | | Feces composition of TS1_V2_turnbaugh of MetaHIT Consortium (additional members) et al. (2011) (original data Turnbaugh et al. (2009)) |

#### Distribution of the differences between abundances at successive time points

We study the differences between abundances at successive time points of the time series in two ways. First we look at the mean absolute value of the difference between abundances at successive time points $\langle | x(t+\delta t) - x(t) | \rangle$ and second we fit the distribution of the ratios of abundances at successive time points $x(t+\delta t)/x(t)$ with a lognormal. More details about the latter can be found in Methods: Width distribution of ratios of abundances at successive time points. The slope of the mean absolute difference $\langle | x(t+\delta t) - x(t) | \rangle$ as a function of the mean abundance in log-log scale ranges between 0.84 and 0.99 (Figure 8). The order of magnitude of the width of the distribution of the ratios of abundances at successive time points $x(t+\delta t)/x(t)$ is one and there is no correlation between the mean abundance and the width of this distribution (Figure 9). The goodness of the fit is characterized by the p-value, a higher p-value corresponds to a better fit of the distribution to a lognormal function (see Methods: Width distribution of ratios of abundances at successive time points). The p-value is represented by the color of the points.

#### Neutrality test

Both neutrality tests, the Kullback-Leibler divergence and the p-value of the neutral covariance test, show that most of the experimental time series are in the niche regime (Figure 10).

#### Noise color

The noise color is predominantly in the pink to white region. It is also independent of the mean abundance (Figure 11).

### Supporting results

#### The noise color depends on the product of the mean abundance and the self-interactions.

To study the noise color, we represented it as a function of different variables. We concluded that the noise color depends on the mean abundance times the self-interaction (Figure 12). This curve can be fitted by a

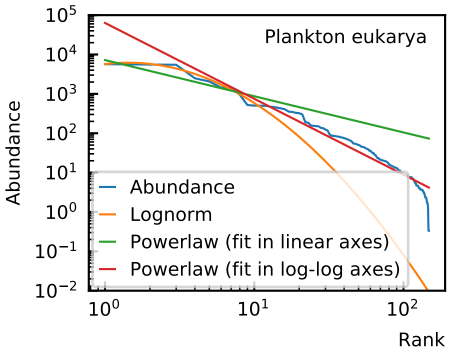

**Figure 7.** The rank abundance curve of experimental data is heavy-tailed. It cannot be exactly fitted by a power law or lognormal distribution.

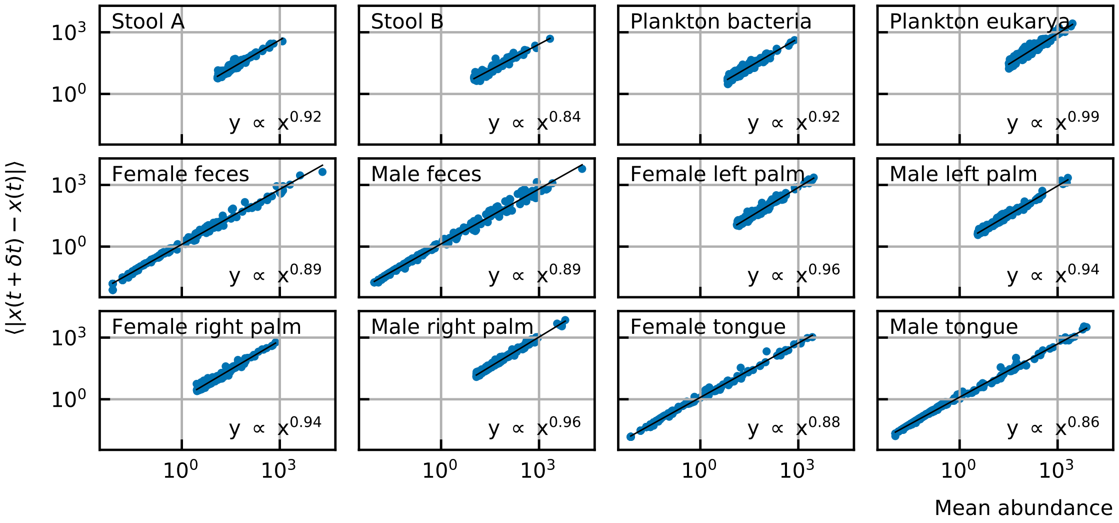

**Figure 8.** For experimental time series, the slope of the mean absolute difference between abundances at successive time points as a function of the mean abundance in log-log scale ranges between 0.84 and 0.99.

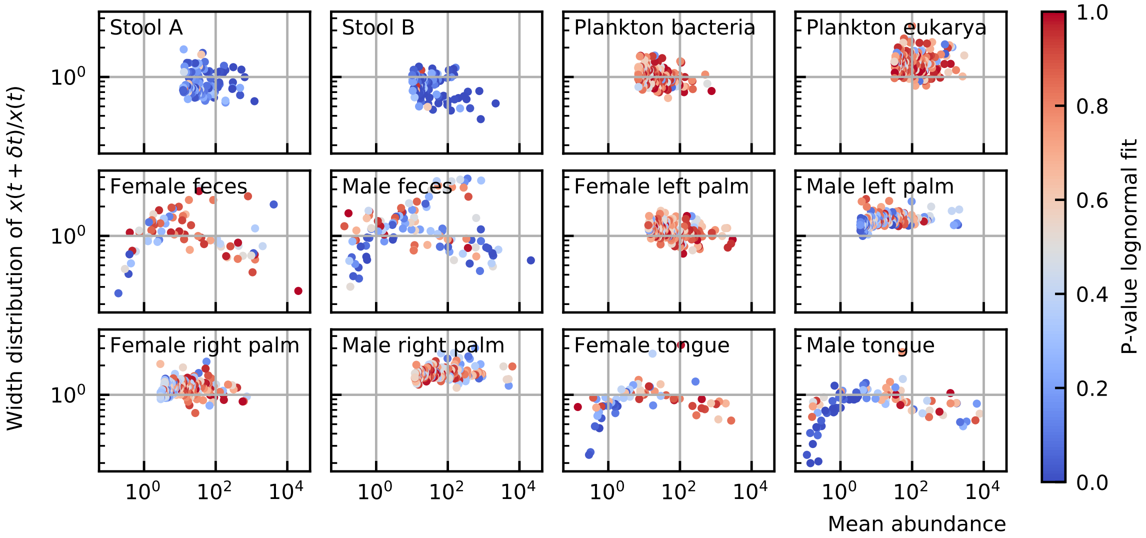

**Figure 9.** For experimental time series, the width of the distribution of the ratios of abundances at successive time points $x(t + \delta t)/x(t)$ is in the order of 1, which means that the fluctuations are large.

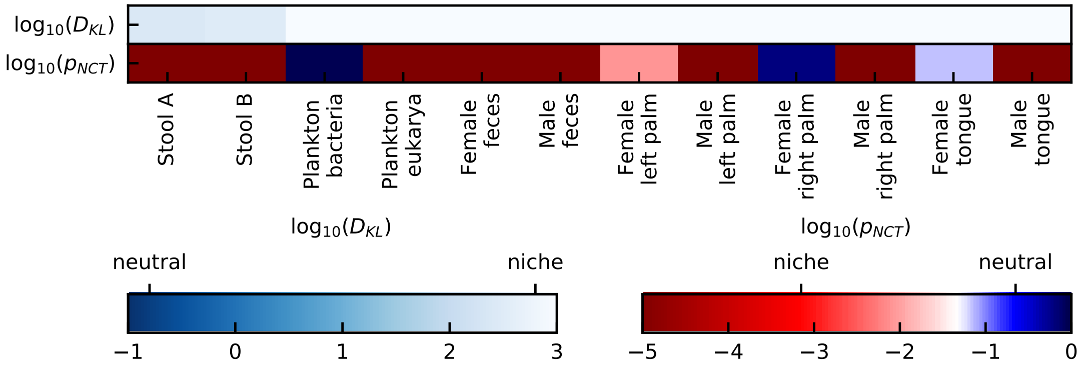

**Figure 10.** Both neutrality tests, the Kullback-Leibler divergence and the p-value of the neutral covariance test, show that most of the experimental time series are in the niche regime.

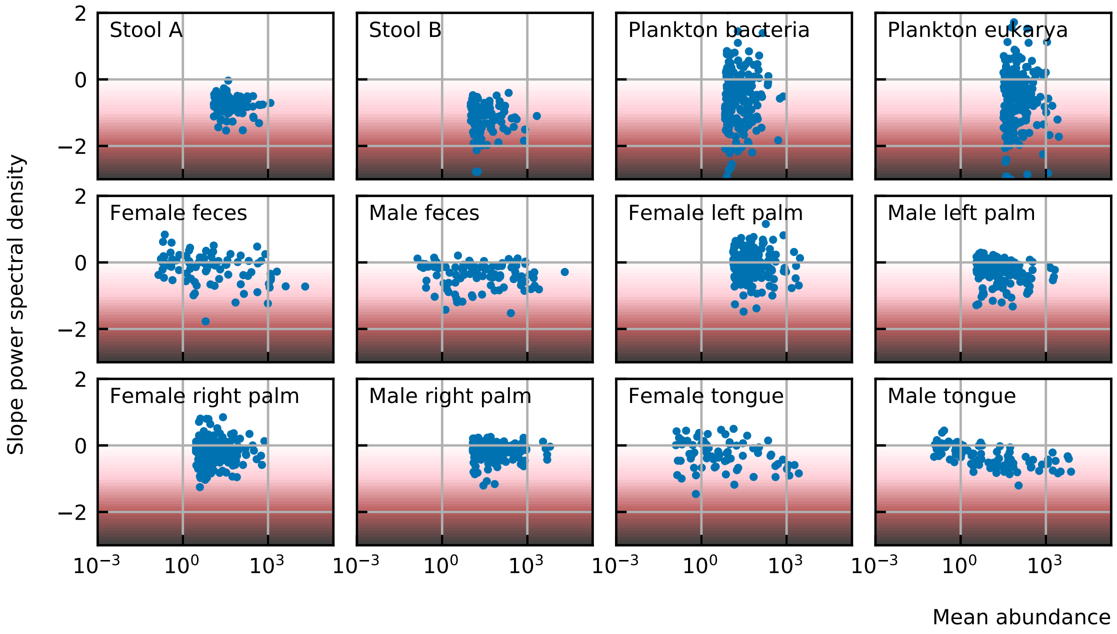

**Figure 11.** For experimental time series, the noise color is predominantly in the pink to white region. It is also independent of the mean abundance.

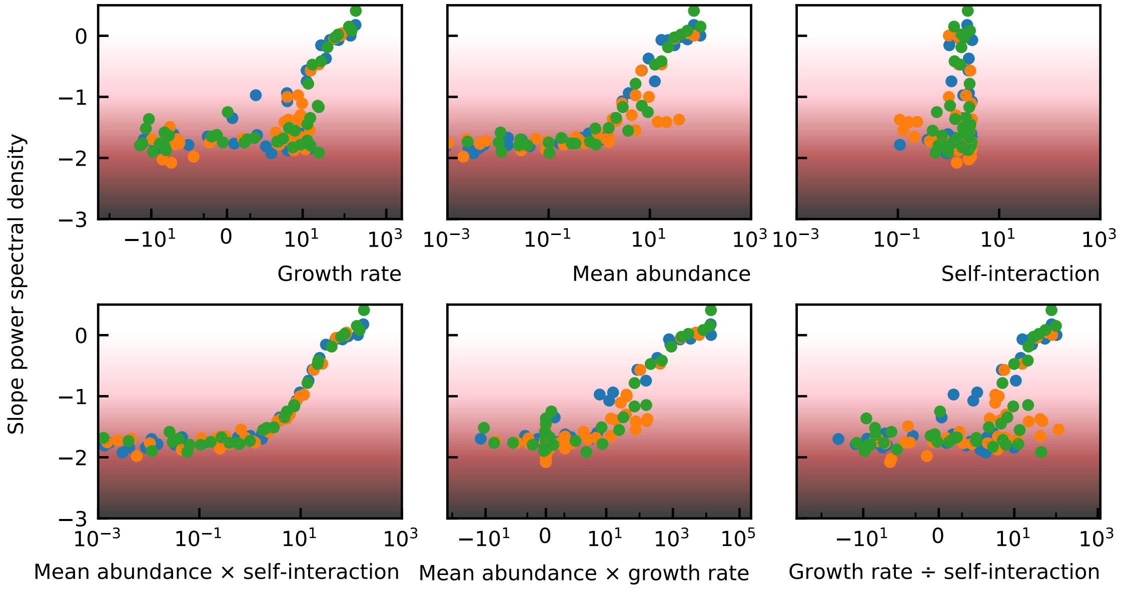

**Figure 12.** The noise color depends on the product of the mean abundance and the self-interaction. The correspondence with the growth rate or mean abundance are less pronounced.

---

sigmoid with $x$-axis in log scale (Figure 13) and the fitted function can be used to calculate the self-interaction of a species given its noise color and mean abundance.

#### All noise characteristics can be obtained with the logistic model.

Given the abundance and the slope of the power spectral density (noise color), we can determine the self-interaction and growth rate of a species via the fitted sigmoid curve (Figure 14). The fitted sigmoid curve gives the value of the product of the mean abundance and self-interaction given the slope of the power spectral density (Figure 14B). Using this value and the relation between the self-interaction and the product of the self-interaction and mean abundance (colored lines of Figure 14F) the self-interaction can be calculated (Figure 14E). For noninteracting species the growth rate equals the product of the mean abundance and self-interaction (black line in Figure 14H), therefore the growth rate is retrieved immediately (Figure 14G).

Once all the parameters are determined, we can perform simulations with large linear multiplicative noise which have the same characteristics as experimental time series. These results are presented in Figure 4 of the main paper.

---

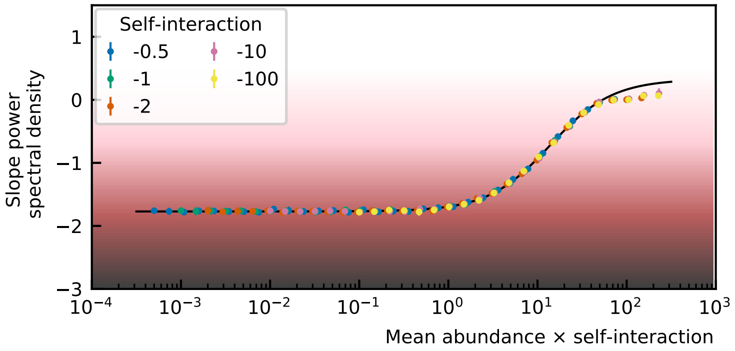

**Figure 13.** The curve of the noise color as a function of the logarithm of the mean abundance and self-interaction can be fitted with a sigmoid function.

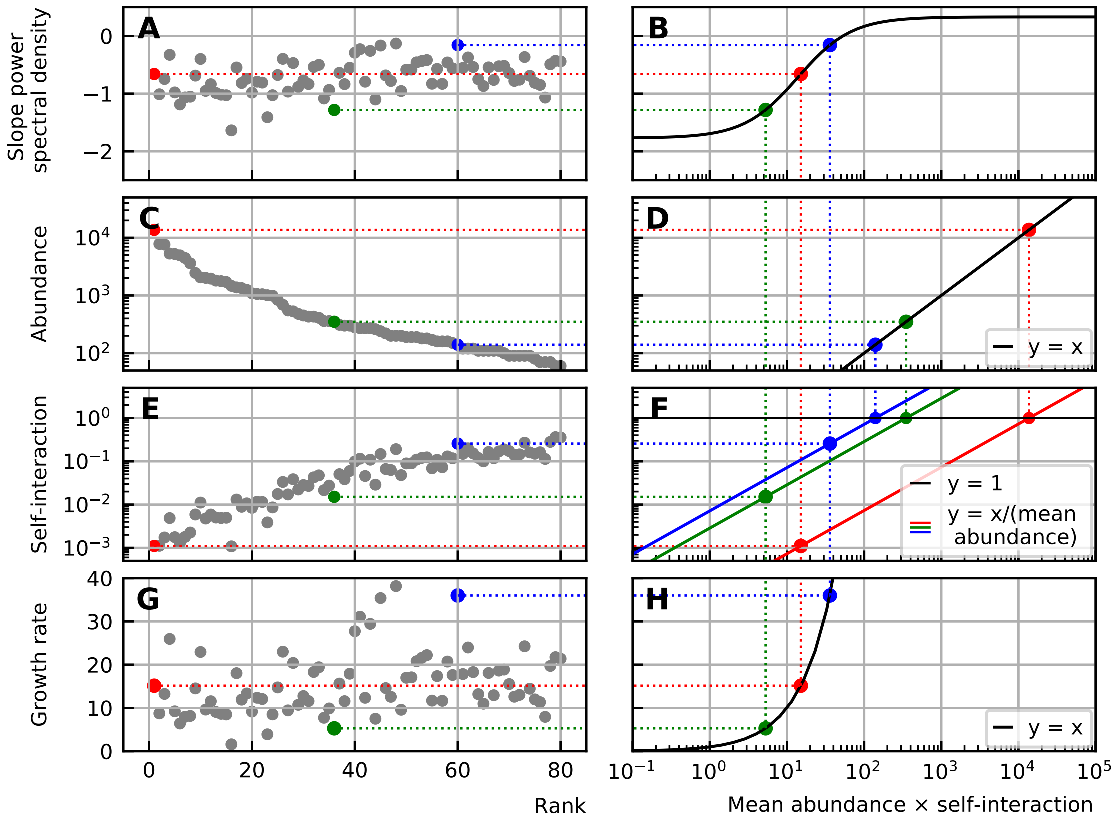

**Figure 14.** This scheme represents how the self-interaction and growth rate can be retrieved from the noise color and abundance data. The fitted sigmoid curve gives the relation between the noise color, abundance and self-interaction (B). Given the abundance (D), the relation between the mean abundance and self-interaction can be drawn (colored lines in F). The value of the noise color can then be translated to a value of the self-interaction (A-B-F-E). For noninteracting species, the growth rate equals the product of the mean abundance and self-interaction. The growth rate is thus easily determined (A-B-H-G).

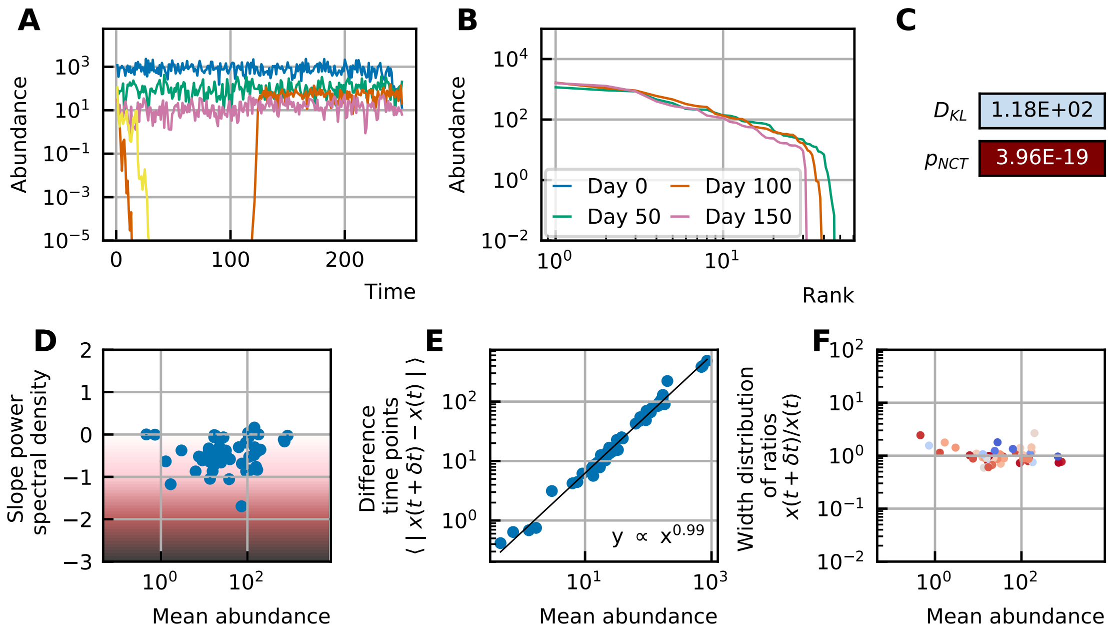

**Figure 15.** All properties of the noise can still be obtained in the presence of interactions: (A) Time series. (B) A heavy-tailed rank abundance that remains stable over time. (C) Results of the neutrality test in the niche regime. (D) Noise color in the white-pink region with no dependence on the mean abundance. (E) The slope of the mean absolute difference between abundances at successive time points is around 1. (F) The width of the distribution of the ratios of abundances at successive time points is in the order of 1 and independent of the mean abundance.

#### The noise characteristics can also be obtained with generalized Lotka-Volterra equations.

In the presence of inter-species interactions, the properties of the experimental time series can also be retrieved. In Figure 15 an example is given with interaction strength 0.02 ($\omega_{ij} \propto \mathcal{N}(0, 0.02)$) and connectance 0.1 which means that 90% of the interactions are set to 0. The rank abundance and self-interactions are derived from the stool A data and the noise is linear with strength $\sigma_{\text{lin}} = 2.5$. The growth rate is calculated given the steady state abundances and interaction matrix. Interactions are however not necessary to obtain the characteristics as seen in the main text.

#### The slope of the differences between abundances at successive time points $x(t + \delta t)/x(t)$ can be fine-tuned by a balanced combination of extrinsic and intrinsic noise.

We can combine extrinsic (linear multiplicative) and intrinsic (square root multiplicative) noise such that the slope of the absolute differences between abundances at successive time points as a function of the mean abundance in log-log space is smaller than one as observed in the experimental time series. For linear multiplicative noise the slope has a value around 1, for sqrt multiplicative noise the value is around 0.66 (Figure 16). A combination where strengths of both noise sources are equal ($\sigma_{\text{lin}} = \sigma_{\text{lin}} = 0.5$) results in an intermediate slope (Figure 16).

#### The width of the distribution of ratios of abundances at successive time points increases with increasing strength of the noise.

The distribution of the ratios of abundances at successive time points $x(t + \delta t)/x(t)$ depends on the rank abundance, self-interaction and interaction strengths. In a first implementation we simulate 50 interacting species with linear noise. The interaction matrix has elements drawn from a normal distribution with mean 0 and standard deviation given by the interaction strength and all self-interactions are imposed to be -1. For increasing noise strength $\sigma_{\text{lin}}$, the width of the distribution of ratios of abundances at successive time points $x(t+\delta t)/x(t)$ increases (Figure 17). In a second implementation, we use 50 species with the rank abundance of the experimental human stool data (Stool A) and self-interactions inferred from the noise color as explained

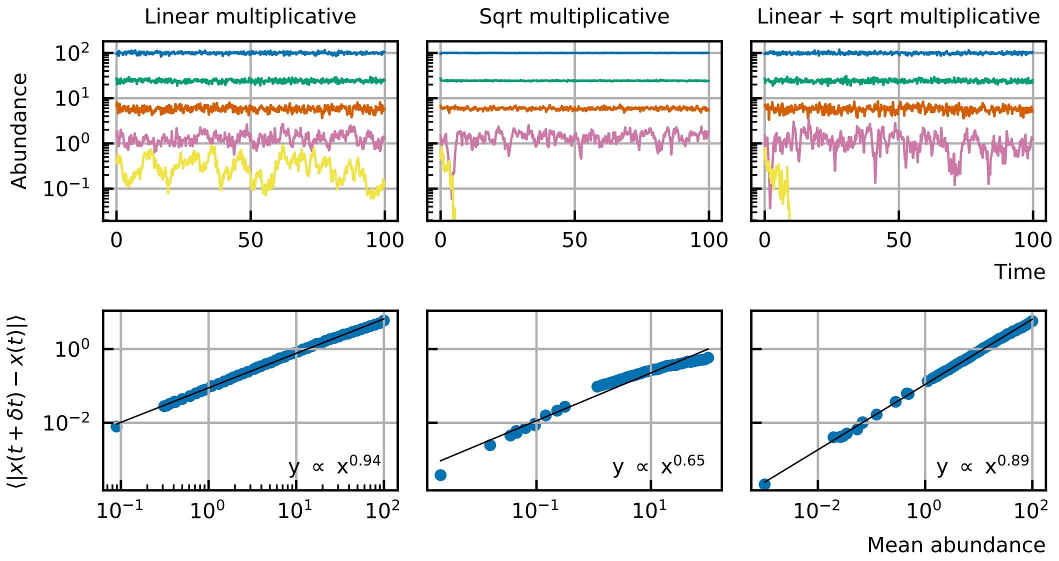

**Figure 16.** The slope of the mean absolute difference between abundances at successive time points with respect to the mean abundance in log-log scale is around 1 (left). For square root noise, this slope is around 0.66 (middle). A mix of linear and square root noise results in intermediate slopes (right). Here, a combination of linear and square root multiplicative noise with equal strengths ($\sigma_{\text{lin}} = \sigma_{\text{sqrt}} = 0.5$) was used to obtain a slope of 0.89.

in the main paper. The interaction matrix for the interacting case is much more sparse here, the connectance is 0.1 which means that only 10% of the interactions are non-zero. The non-zero interactions are distributed like a normal distribution with mean 0 and standard deviation 0.03. As expected, increasing the strength of the noise increases the width of the distribution and adding interaction also slightly increases the width (Figure 18).

### Neutrality of generalized Lotka-Volterra models

The higher the interaction strength the more niche the time series become (Figure 19).

### Rounding time series to integer numbers makes the noise color lighter.

For low abundances there is a finite precision effect. When we calculate the noise color for the time series with a precision up to the integer, we see that the noise color becomes lighter for smaller abundances (Figure 20). This is consistent with the fact that the noise color is correlated with the autocorrelation and the amount of structure in the time series, that is the predictability of the time series. In the time series that have smaller precision, there is less information and the time series is less predictable, therefore the noise color is lighter.

### The self-organized instability model can be reproduced by the stochastic generalized Lotka-Volterra model.

In the main text we discussed that next to stochastic generalized Lotka-Volterra models there exist another algorithm to model interacting species stochastically, the individual based model. We mention in particular the self-organized instability (SOI) model introduced by Solé et al. (2002). The question arises whether the SOI model is equivalent to the generalized Lotka-Volterra models or if it can produce more or different characteristics. An elaborate study of the SOI has been performed in Faust et al. (2018). Most results are in agreement with our findings. However in a supplemental figure of Faust et al. (2018) (S3), we see that the percentage of taxa with pink noise increases and the percentage of taxa with brown noise decreases when the stochasticity increases, where the stochasticity is defined as the ratio between the mean of extinctions

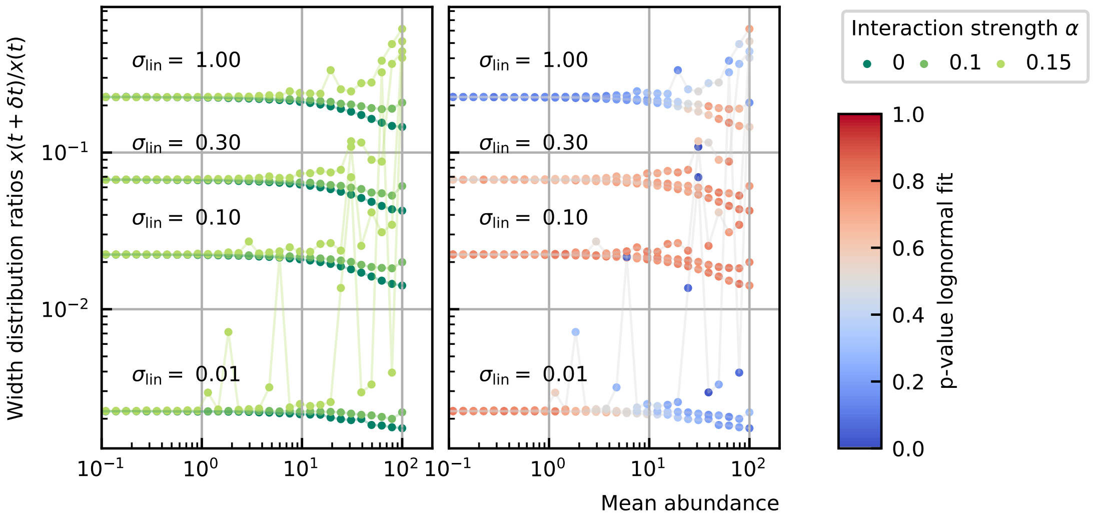

**Figure 17.** The width of the distribution of the ratios of abundances at successive time points $x(t + \delta t)/x(t)$ for time series of 50 interacting species with equal abundances increases with increasing strength of the linear noise. For high abundances, the width also increases with the interaction strength.

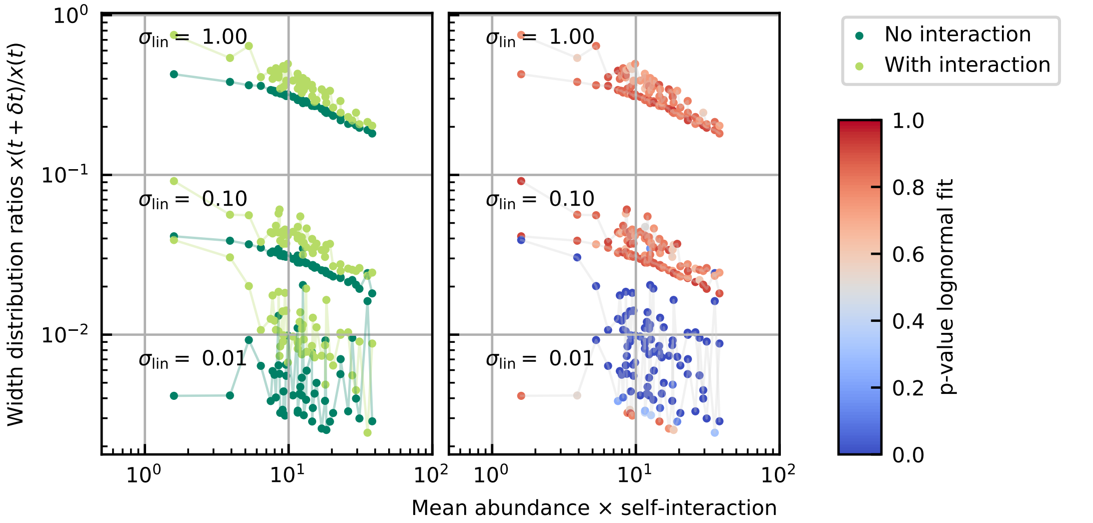

**Figure 18.** The width of the distribution of the ratios of successive time points for time series for 50 interacting species with a rank abundance and inferred self-interaction of the stool A data. The width increases with increasing strength of the linear noise.

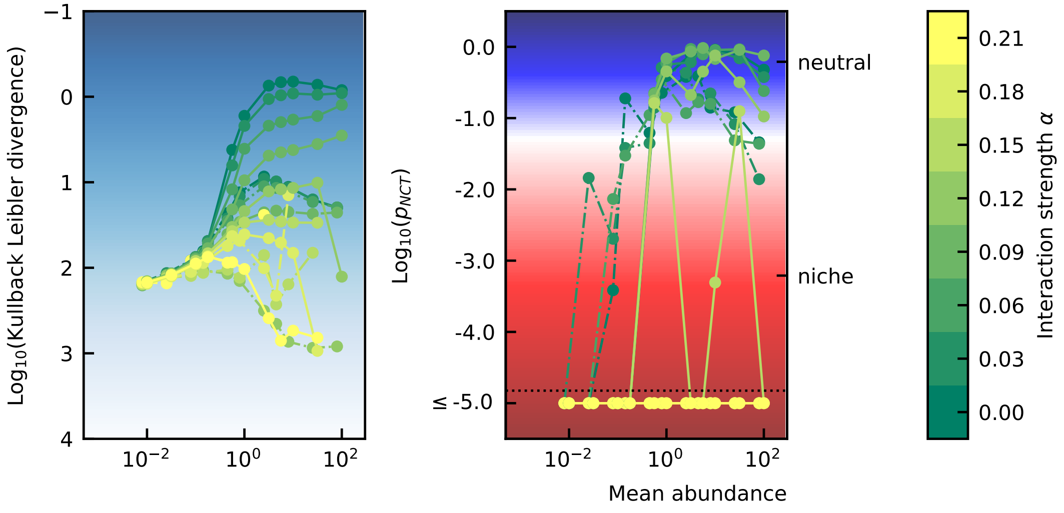

**Figure 19.** The higher the interaction strength the more niche the time series become.

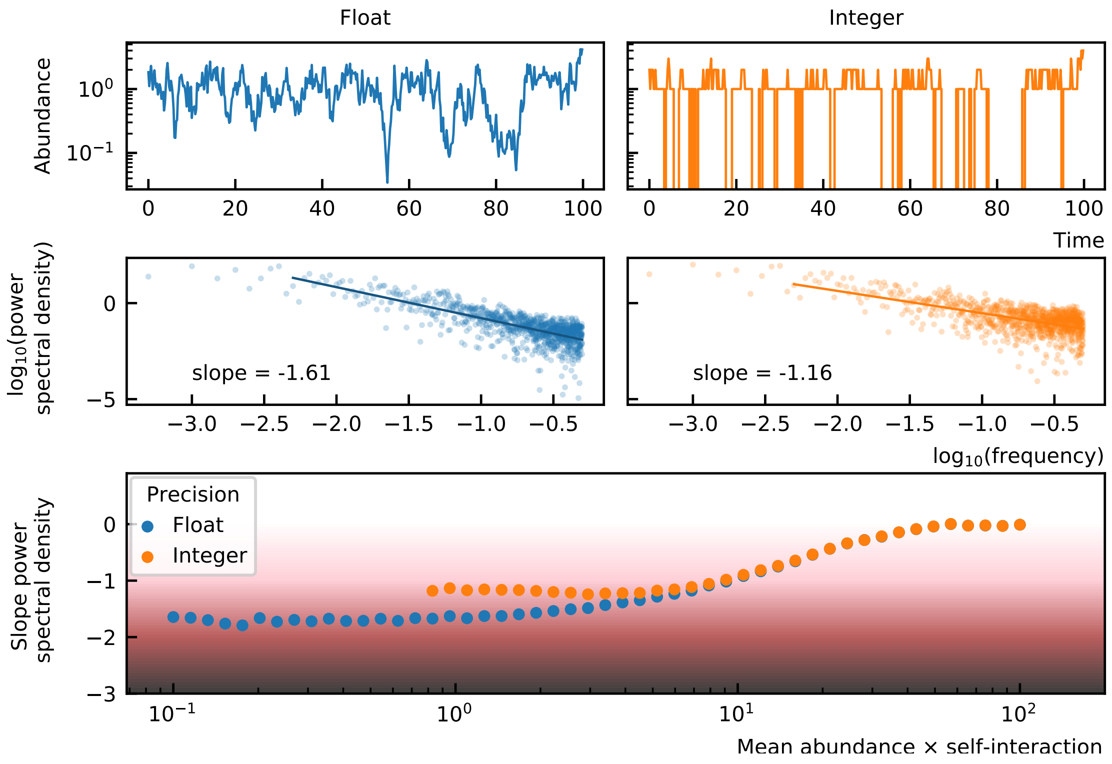

**Figure 20.** Finite precision makes the noise color lighter for species with small abundances.

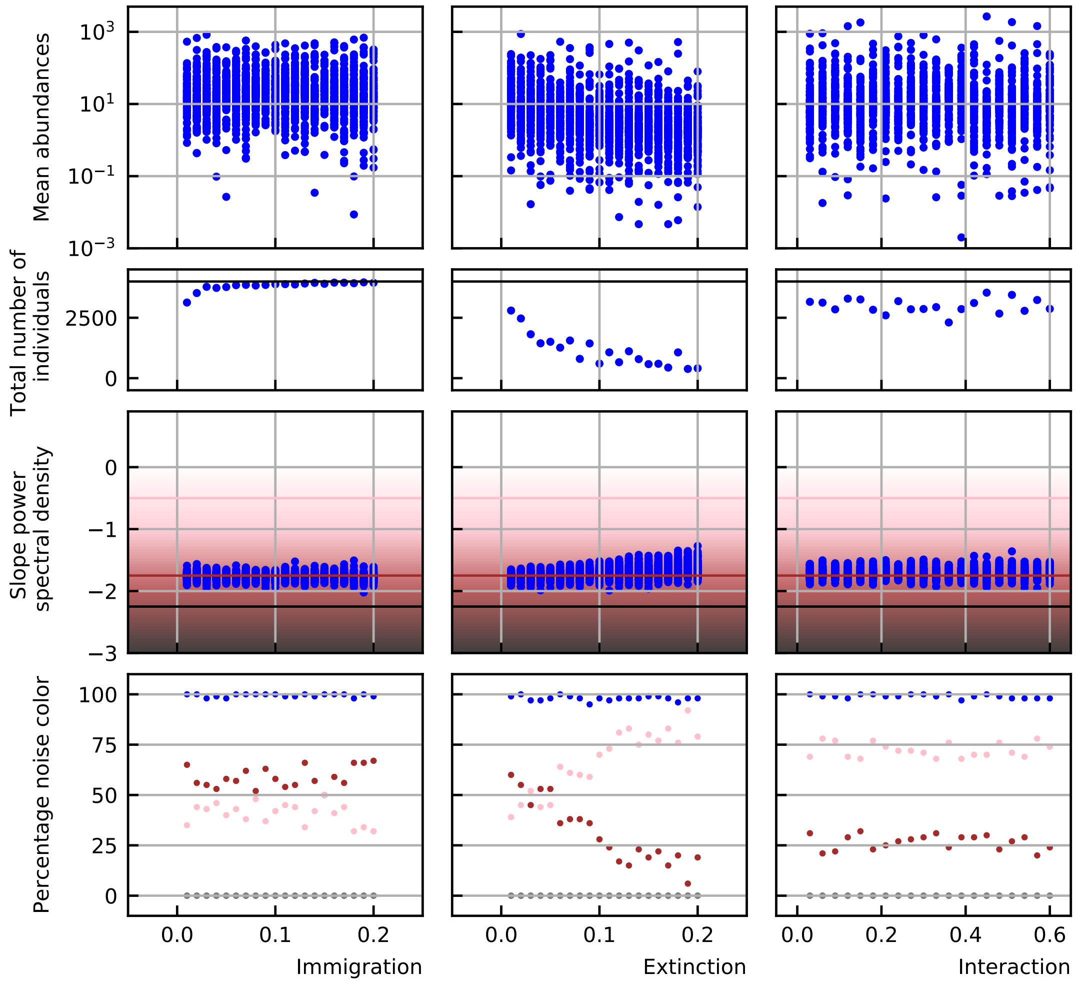

**Figure 21.** Time series simulations were performed for multiple values of the immigration, extinction and interaction rate. Shown are the mean abundance values of these simulations (top panels), the total number of individuals (second row), the distribution of the noise colors (third row) and the percentages of all noise colors as delimited by Faust et al. (2018) (bottom row). For the SOI model, an increase in the extinction rate causes a shift from brown to pink noise. At the same time the mean abundances and total number of individuals decrease.

---

and immigrations and the mean of the absolute interaction strengths (excluding diagonal values). This cannot be easily concluded from our previous results and we therefore studied this aspect in more detail.

We first evaluated the contributions of the different parameters. We therefore run simulations using the code provided by the authors of Faust et al. (2018), only changing one of the parameters at a time. What we notice is that the shift from brown to pink noise is mostly caused by the extinction rate and not the immigration rate or interaction strength (Figure 21, parameters are in Table 2 and Table 3).

In Figure 21, we also see that with increasing extinction the mean abundance is decreasing (top panel). At the same time the noise color is becoming lighter (shifting to pink). To understand the connection between the parameters, we made a figure with both the noise color and mean abundance on the axes for two time series (Figure 22A). We see that the noise color becomes more pink for smaller mean abundances. How do these results relate to the results for the generalized Lotka-Volterra equations?

There are multiple differences between the SOI and gLV models. First of all, the SOI model follows an individual based approach. It is based on the Gillespie algorithm with K-leap method. At every time step the propensities for all events are calculated and K processes are chosen accordingly. The elapsed time

**Table 2.** Parameter values of varied parameters.

| Column in figure 1 | 1 | 2 | 3 |
|---|---|---|---|
| Varying parameter | Immigration rate | Extinction rate | Interaction strength |
| Immigration (m) | $\mathcal{U}$(0, immigration) | $\mathcal{U}$(0, 0.02) | $\mathcal{U}$(0, 0.1) |
| Extinction (e) | $\mathcal{U}$(0, 0.01) | $\mathcal{U}$(0, extinction) | $\mathcal{U}$(0, 0.1) |
| Interaction strength (A) | $\mathcal{U}$(-0.2, 0.2) | $\mathcal{U}$(-0.2, 0.2) | $\mathcal{U}$(-interaction, interaction) |

**Table 3.** Parameter values of fixed parameters.

| Parameter | Value |
|---|---|
| Number of individuals / lattice sites (I) | 4000 |
| Number of species (S) | 100 |
| Positive edge percentage (PEP) | 20 |
| Diagonal elements | -0.5 |
| Connectance | 0.02 |

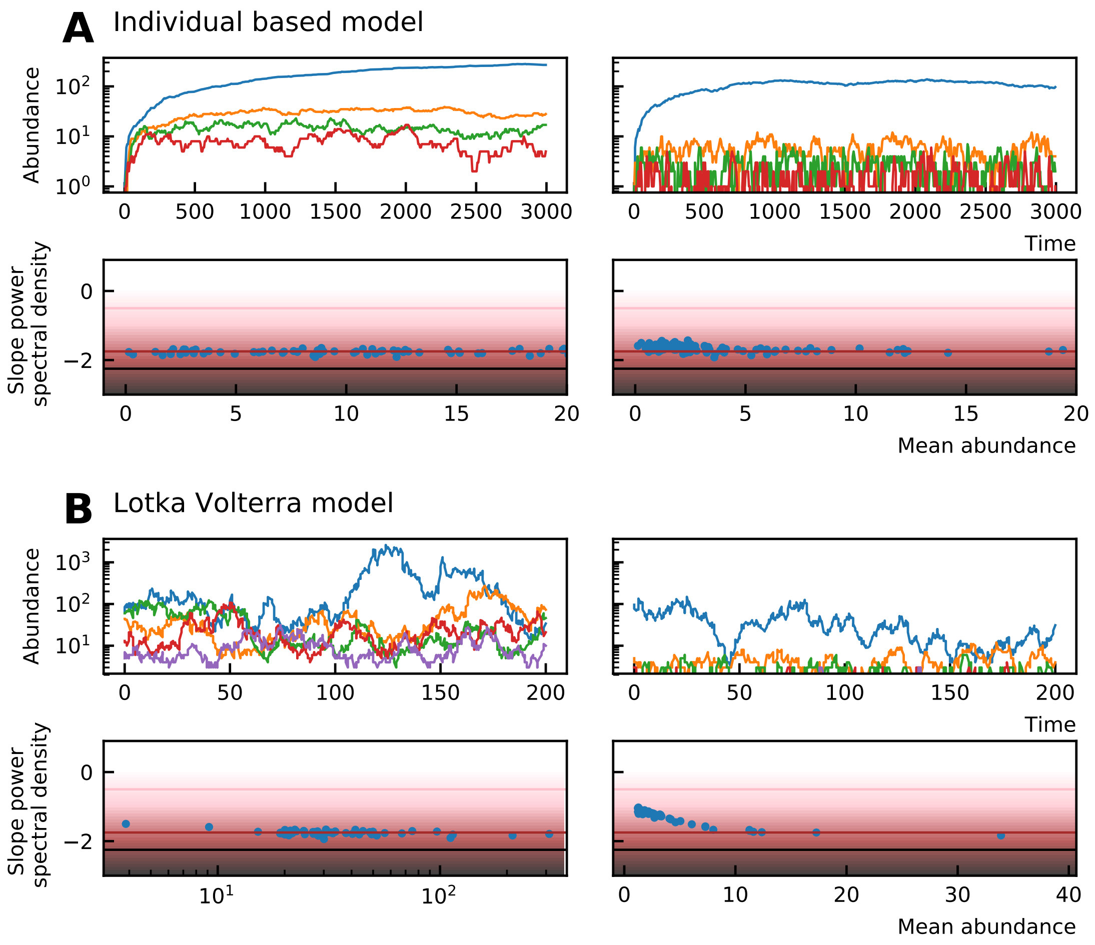

**Figure 22.** (A) Time series generated by the SOI model for different extinction rates, $e = 0.01$ for the left column and $e = 0.16$ for the right column (additional parameters can be found in the second column of Table 2 and Table 3). The lower figures show the noise color as a function of the mean abundance for the corresponding time series. For higher extinction (right side) the mean abundances are lower and the noise color becomes lighter with decreasing mean abundance. (B) Time series generated with the gLV model and rounded to integer precision after the simulation for different extinction rates, $e = 0.001$ on the left and $e = 0.8$ on the right. The noise color becomes lighter for low abundances.

$\tau$ is calculated by the gamma function. Since it is an individual based model, the abundances will only take integer numbers. GLV models are continuous and the abundances of the time series are real positive numbers. We showed in the previous section that Rounding time series to integer numbers makes the noise color lighter.

A second difference between the SOI and our gLV model is that the SOI model is based on immigration and that no species can grow in the absence of other species. In the gLV model, we do not consider immigration and the growth rates of species are often positive. Furthermore the interaction matrix is interpreted in a different way. Given the interaction matrix used for the SOI model ($\omega$), the interaction matrix of the gLV model ($\omega'$) needs to obey the following rules for the time series to be comparable:

1. $\omega'_{ii} = 0$ all self-interactions are zero,
2. if $\omega_{ij} > \omega_{ji} > 0 : \omega'_{ij} = \omega_{ij} + \omega_{ji}$ and $\omega'_{ji} = 0$ mutualistic interactions are transformed into commensalistic interactions and,
3. if $\omega_{ij} < 0$ and $\omega_{ij} < \omega_{ji} : \omega'_{ij} = \omega_{ji} - \omega_{ij}$ and $\omega'_{ji} = 0$ competitive / parasitic interactions are transformed into amensalistic interactions.

The most important difference is that the self-interactions are zero. For gLV models the self-interaction needs to be negative such that the abundance remains bounded and cannot go to infinity. In the SOI model this constraint is imposed by the maximal number of individuals. The propensity for growth decreases linearly with the number of individuals. When the maximal number of individuals is attained all growth propensities are zero.

If we include all the aforementioned elements:

1. add immigration,
2. consider only negative growth rates (extinction),
3. small self-interaction,
4. round the results to integer values after performing the complete time series,

then we obtain similar results as the SOI model (Figure 22B and Figure 23).

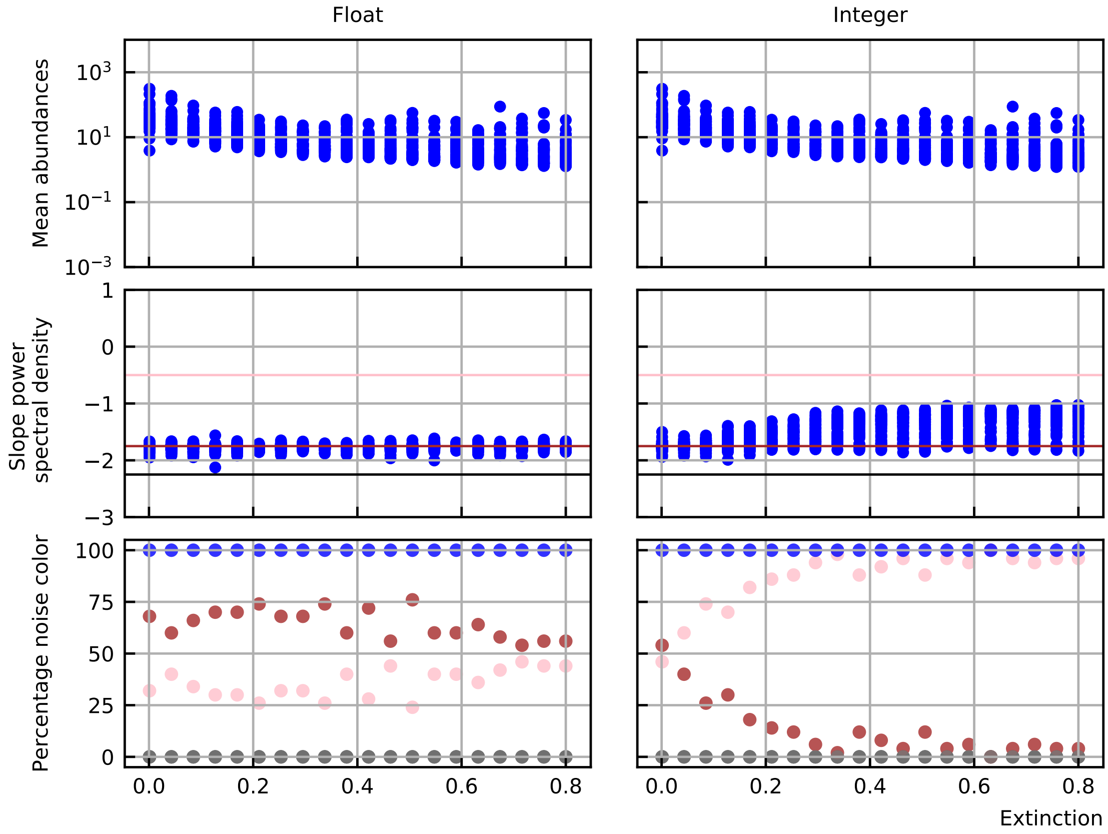

**Figure 23.** When imposing different constraints —addition of immigration, negative growth rates, small self-interaction, small precision— on the gLV model, the noise color shifts for increasing extinction as in the SOI model.

**MetaHIT Consortium (additional members)**, Arumugam M, Raes J, Pelletier E, Le Paslier D, Yamada T, Mende DR, Fernandes GR, Tap J, Bruls T, Batto JM, Bertalan M, Borruel N, Casellas F, Fernandez L, Gautier L, Hansen T, Hattori M, Hayashi T, Kleerebezem M, et al. Enterotypes of the human gut microbiome. Nature. 2011 May; 473(7346):174–180. http://www.nature.com/articles/nature09944, doi: 10.1038/nature09944.

**Solé RV**, Alonso D, McKane A. Self-organized instability in complex ecosystems. Philosophical Transactions of the Royal Society of London Series B, Biological Sciences. 2002 May; 357(1421):667–671. doi: 10.1098/rstb.2001.0992.

**Turnbaugh PJ**, Hamady M, Yatsunenko T, Cantarel BL, Duncan A, Ley RE, Sogin ML, Jones WJ, Roe BA, Affourtit JP, Egholm M, Henrissat B, Heath AC, Knight R, Gordon JI. A core gut microbiome in obese and lean twins. Nature. 2009 Jan; 457(7228):480–484. http://www.nature.com/articles/nature07540, doi: 10.1038/nature07540.

**Walczak AM**, Mugler A, Wiggins CH. Analytic Methods for Modeling Stochastic Regulatory Networks. In: Liu X, Betterton MD, editors. *Computational Modeling of Signaling Networks*, vol. 880 Totowa, NJ: Humana Press; 2012.p. 273–322. http://link.springer.com/10.1007/978-1-61779-833-7_13, doi: 10.1007/978-1-61779-833-7_13.

**Washburne AD**, Burby JW, Lacker D. Novel Covariance-Based Neutrality Test of Time-Series Data Reveals Asymmetries in Ecological and Economic Systems. PLOS Computational Biology. 2016 Sep; 12(9):e1005124. https://dx.plos.org/10.1371/journal.pcbi.1005124, doi: 10.1371/journal.pcbi.1005124.

**Wennekes PL**, Rosindell J, Etienne RS. The Neutral—Niche Debate: A Philosophical Perspective. Acta Biotheoretica. 2012 Sep; 60(3):257–271. http://link.springer.com/10.1007/s10441-012-9144-6, doi: 10.1007/s10441-012-9144-6.

**Zhu C**, Yin G. On competitive Lotka–Volterra model in random environments. Journal of Mathematical Analysis and Applications. 2009 Sep; 357(1):154–170. https://linkinghub.elsevier.com/retrieve/pii/S0022247X09002777, doi: 10.1016/j.jmaa.2009.03.066.
